# Dicarboxylic acid supplementation protects from acute kidney injury via stimulation of renal peroxisomal activity

**DOI:** 10.1101/2023.05.31.543068

**Authors:** Anne C. S. Barbosa, Katherine E. Pfister, Takuto Chiba, Joanna Bons, Jacob P. Rose, Jordan B. Burton, Christina D. King, Amy O’Broin, Victoria Young, Bob Zhang, Bharathi Sivakama, Alexandra V. Schmidt, Rebecca Uhlean, Birgit Schilling, Eric S. Goetzman, Sunder Sims-Lucas

## Abstract

**Introduction:** Lysine succinylation is a post-translational modification associated with the control of several diseases, including acute kidney injury (AKI). It is suggested that hypersuccinylation favors peroxisomal fatty acid oxidation (FAO) instead of mitochondrial. In addition, the medium-chain fatty acids (MCFAs) dodecanedioic acid (DC_12_) and octanedioic acid (DC_8_), upon FAO, generate succinyl-CoA, resulting in hypersuccinylation. DC_8_ is convenient, inexpensive, easily administered, and efficient. We believe this study could be translated in the future to clinical settings, which would highly benefit patients at high risk of AKI.

**Methods and Results:** To test the protective roles of MCFAs during AKI, mice were fed with control, 10% DC_12_, or 10% DC_8_ diet, then, subjected to either ischemic-AKI, or cisplatin-AKI models. Supplementation was provided until sacrifice. Biochemical, histologic, genetic, and proteomic analysis were performed, the latter involving a lysine-succinylome-based analysis. Both DC_8_ and DC_12_ prevented the rise of AKI markers in mice that underwent renal injury. However, DC_8_ was even more protective against AKI than DC_12_. Finally, succinylome analysis evidenced that the kidneys of DC_8_-fed mice showed an extensive succinylation of peroxisomal activity-related proteins, and a decline in mitochondrial FAO, in comparison to control-fed mice.

**Conclusion:** DC_8_ supplementation drives renal protein hypersuccinylation, promoting a shift from mitochondrial to peroxisomal FAO, and protecting against AKI.

**Significance Statement:** Lysine succinylation of proteins is shown to control several diseases, including acute kidney injury (AKI). Here we show that mice supplemented with the medium-chain fatty acid octanedioic acid successfully presented a high level of succinylation and were protected from both ischemia-reperfusion- and cisplatin-induced AKI. Moreover, our study demonstrates that peroxisomal activity was increased while mitochondrial activity was preserved, suggesting that the metabolism of diet-obtained medium-chain fatty acids by peroxisomes is renoprotective.

## Introduction

Acute kidney injury (AKI) is associated with high morbidity and mortality worldwide and no specific treatment for AKI has been clinically approved thus far. The major causes of AKI are ischemia, due to physical trauma; as well as sepsis, major cardiac or vascular surgeries and nephrotoxins, such as the widely used chemotherapy drug cisplatin [1, 2].

Proximal renal tubule cells are the most abundant cells present in the kidney cortex and they require adenosine triphosphate (ATP) as a fuel, which is efficiently obtained via β-oxidation of fatty acids. Fatty acid oxidation (FAO) takes place primarily in mitochondria. However, the peroxisomes are another metabolic organelle with a parallel FAO pathway to the mitochondrial pathway [3–5]. During ischemic AKI, the sudden loss of oxygen damages mitochondria as well as peroxisomes, and renal FAO becomes typically impaired for several days post-injury [6, 7]. This likely exacerbates injury and delays kidney repair, with consequences for the whole body. In addition to abundant mitochondria, proximal tubules are rich in peroxisomes. While peroxisomes do not produce energy directly, recent evidence indicates they transfer short-chain FAO products, such as propionyl-CoA and carnitine ester, to the mitochondria that can subsequently contribute to energy production in the mitochondria [8–13].

Protein post-translational modifications (PTMs) are evolutionarily conserved mechanisms for regulating cellular homeostasis. In the past decade, lysine succinylation, which arises from proteins chemically reacting with the tricarboxylic acid (TCA) cycle metabolite succinyl-CoA [14], was unveiled as a PTM associated with important physiological functions [5]. In addition, sirtuin 5 (Sirt5) has been identified as a lysine deacylase capable of reversing lysine succinylation [15, 16]. Mice lacking Sirt5 exhibit hypersuccinylation of many enzymes involved in β-oxidation- as well as ketogenesis-related pathways [15]. This hypersuccinylation is believed to suppress function, as evidenced by accumulation of medium- and long-chain acylcarnitines, which are intermediates of intracellular fatty acid metabolism [12, 17]. Sirt5 is localized to both mitochondria and peroxisomes, and we have previously shown that upon renal ischemia-reperfusion (IRI) or cisplatin treatment, Sirt5 knockout mice are protected against AKI [7, 15, 16]. These mutants also presented augmented peroxisomal FAO whereas their mitochondrial FAO function was diminished, indicating Sirt5 plays an important role in balancing mitochondrial and peroxisomal FAO in renal tubule cells [7]. Moreover, there is increasing evidence that bolstering peroxisomal FAO can protect against AKI. For example, inhibiting mitochondrial FAO with etomoxir leads to increased compensatory peroxisomal FAO and protection against AKI [18].

While the above studies demonstrate proof-of-concept that boosting kidney peroxisomal FAO protects against AKI, it remains challenging to manipulate the peroxisomal pathway therapeutically. Inhibitors like etomoxir are associated with serious side effects due to inhibition of FAO in liver and other tissues [19]. Here, our approach was to feed mice a diet enriched with dicarboxylic fatty acids (DCAs), which are believed to be predominantly metabolized by peroxisomes rather than mitochondria [10, 20, 21]. DCAs are not found in the diet. Rather, they are produced by liver and kidney from their monocarboxylic counterparts, most notably 12-carbon and 10-carbon chain lengths (C10 and C12), by a pathway known as fatty acid ω-oxidation, occurring in the endoplasmic reticulum and cytoplasm. This pathway becomes active in the liver during fasting and during times of heavy medium-chain fatty acid ingestion, such as what occurs with medium-chain triglyceride (MCT) therapy. By enriching the diet with DCAs, we bypass the ω-oxidation pathway, and consequently show that the resulting drive on peroxisomal FAO protects against multiple forms of AKI in mice.

## Materials and Methods

### Animals and dicarboxylic acid supplementation

Wild-type mice were obtained from the Jackson Laboratory (B6129SF1/J, stock no.101043 for renal IRI, 129S1/SvImJ, stock no. 002448 for cisplatin-AKI). Age-matched 9- to 14-week-old male mice were used throughout the study. The University of Pittsburgh Institutional Animal Care and Use Committee approved all experiments (approval no. **22112009**).

Animals were fed *ad libitum* primarily for seven days prior to and seven days after IRI; or seven days prior to and three days after an in-bolus cisplatin injection. For the wash in wash out experiments, mice were fed 10% DC_8_ for 1, 3, and 5 days followed by a wash out of 1, 3 and 5 days. Groups were divided into control diet, 10% DC_12_ (Sigma-Aldrich D1009) w/w, or 2.5-10% DC_8_ (Alfa Aesar A13963) w/w supplementation in their control mush diet.

### Ischemia-reperfusion injury (IRI) AKI Model

Ischemic AKI was induced by a unilateral renal ischemia-reperfusion injury (IRI) with a delayed contralateral nephrectomy as previously described [7, 22, 23]. Briefly, mice were anesthetized with 2% isoflurane. Core body temperature of the mice was monitored with a rectal thermometer probe and was maintained at 37–37.5°C throughout the procedures with a water-heating circulation pump system (EZ-7150; BrainTree Scientific) and an infrared heat lamp (Shat-R-Shield). Ethiqa®, extended-release buprenorphine 1.3 mg/mL (Fidelis MIF 900-014), was administered for analgesia (3.25 mg/kg body weight, administered subcutaneously) as a single dose. With aseptic techniques, a dorsal incision was made to expose the kidney, and renal ischemia for 26 minutes was induced by clamping of the left kidney pedicle with a nontraumatic microvascular clamp (18055–04; Fine Science Tools). Renal reperfusion was visually verified afterwards. Contralateral nephrectomy of the right kidney was performed at day 6. Mice were euthanized at day 7 to harvest blood and the injured kidney. Serum was separated from the blood and analyzed by the Kansas State Veterinary Diagnostic Laboratory for levels of creatinine, BUN, and other biochemical markers.

### Cisplatin nephrotoxic AKI Model

To induce cisplatin AKI, mice were given a single dose of 20 mg/kg intraperitoneal cisplatin (Fresenius Kabi), or vehicle control of normal saline as described previously [7]. Mice were euthanized on day 3 to harvest blood and the kidneys.

### Western blotting

Kidney tissues were lysed in radioimmunoprecipitation assay buffer (Thermo Fisher Scientific), and the homogenates plated in triplicate to measure protein concentration using a Bradford assay kit (Bio-Rad). The following antibodies were used in this study: anti-NGAL (R&D systems AF1857; Abcam ab702p7), anti-PEX5 (CST 830205), anti-pmp70 (Abcam ab85550), anti-Thioredoxin (Sigma SAB5300166), Anti-Succinyllysine (PTM Bidabs Ptm-419), 8-OHdg (Bioss bs-1278R), Catalase (Invitrogen PAS-88250). Anti-GAPDH or anti–a/b-tubulin were used as loading controls.

### Real-Time PCR

Real-time PCR analysis was performed as previously described to determine mRNA level in whole kidneys [7]. Complementary DNA was reverse-transcribed from 500 ng of total RNA with SuperScript First-Strand Synthesis System II (Thermo Fisher Scientific). Real-time PCR analysis was performed with gene specific primer oligos, SsoAdvanced SYBR Green Super-mix (Bio-Rad), and CFX96 Touch Real-Time PCR Detection System with C1000 Thermal Cycler (Bio-Rad). Cycling conditions were 95°C for 10 minutes, then 40 cycles of 95°C for 15 seconds and 60°C for 1 minute. Rn18S was used for endogenous control (Primer sequence forward AGAAACGGCTACCACATCCA; primer sequence reverse TACAGGGCCTCGAAAGAGTC). NGAL expression (Primer sequence forward GCAGGTGGTACGTTGTGGG; primer sequence reverse CTCTTGTAGCTCATAGATGGTGC) was compared with the Rn18S endogenous control and analyzed using the 2_2ΔΔCt_ method.

### Tissue Section Analysis

Kidneys were fixed in 4% paraformaldehyde and embedded in paraffin. These tissues were sectioned at 4 m. The kidney sections were stained with hematoxylin and eosin (H&E). H&E stained slides were subjected to histologic evaluation, and 340 magnification images of renal cortex and medulla were obtained with a Leica DM2500 optical microscope. Semiquantitative scoring (0– 4) for tubular injury was performed in blind fashion in terms of tubular dilation, proteinaceous cast formation, and loss of brush border. Immunostaining were performed as previously described [7, 22], with paraffin-embedded tissues and with following primary antibodies: anti-NGAL (R&D systems AF1857; Abcam ab702p7), anti-PEX5 (CST 830205), anti-pmp70 (Abcam ab85550), anti Kidney Injury Marker 1 (R&D Biosystems MAB1817). Antibodies were used at 1:50 to 1:200. Followed by conjugation with ALEXA^®^ fluorescent antibodies at 1:200-1:500.

### TUNEL assay

Terminal deoxynucleotidyl transferase-mediated digoxigenin-deoxyuridine nick-end labeling (TUNEL) staining was performed by using the In Situ Cell Death Detection Kit from Roche (Mannheim, Germany). Briefly, paraffin-embedded tissues were deparaffinized, antigen-retrieved, and incubated with TUNEL as previously described [24].

### Quantitative Mass Spectrometry

Both 7-day renal IRI and contralateral kidney tissues from control- (n=4), 10% DC_12_- (n=3), and 10% DC_8_-fed (n=4) were homogenized, trypsinized, and then succinylated peptides were enriched using the PTMScan Succinyl-Lysine Motif Kit (Cell Signaling Technologies) as previously described but with slight adaptations (also see Supplemental Appendix 1) [7]. Using reversed-phase high performance liquid chromatography/electrospray ionization-tandem mass spectrometry (HPLC/ESI-MS/ MS), succinyl-enriched samples were analyzed by data-independent acquisition (DIA) on an Orbitrap Eclipse mass spectrometer [25–27] and site-specific changes in succinylation were quantified using library-free directDIA in Spectronaut (Biognosys) [25, 28]. Parallel whole lysate LC-MS/MS analysis conducted on an Orbitrap Exploris 480 mass spectrometer operated by DIA-MS was used for PTM-site normalization adjusting for the protein-level changes. See further experimental and statistical details in Supplemental Appendix 1 [27, 29–42].

### Radiolabeled Fatty Acid Oxidation Assays

14C-labeled <u>palmitic acid</u> (C16) and ^14^C-octanoic acid (C_8_) were from PerkinElmer while 14C-DC12 and 14C-lignoceric acid (C_24_) were from Moravek, Inc. All substrates were solubilized in 10 mg/ml α-cyclodextrin and added to a final concentration of 50 µM in reactions containing 100 mM sucrose, 10 mM Tris–HCl, 5 mM KH2PO4, 0.2 mM EDTA, 0.3% fatty acid-free BSA, 80 mM KCl, 1 mM MgCl2, 0.2 mM L-carnitine, 0.1 mM malate, 0.05 mM coenzyme A, 2 mM ATP, 1 mM DTT. Reactions were started with addition of freshly prepared kidney lysate (∼100 µg) and incubated in a rotating water bath at 37°C for 1 hr. For ^14^C-C_16_, which can be metabolized by either mitochondria or peroxisomes, FAO was measured ± 100 µM etomoxir to inhibit the mitochondrial pathway and allow determination of the peroxisomal contribution. Reactions were stopped with perchloric acid, centrifuged to clear precipitated material, and then the water-soluble 14C-labeled FAO products were isolated by extraction with chloroform/methanol and subjected to scintillation counting. For C_8_, which is water-soluble and does not separate from its FAO products upon chloroform/methanol extraction, the rate of FAO was determined by capturing ^14^C-CO_2_ after addition of the perchloric acid. For all substrates, concurrent reactions with no tissue lysate were used as blank controls. Data were normalized to protein concentration.

### Oroboros High-Resolution Respirometry

Freshly prepared kidney homogenates were analyzed with an Oroboros Oxygraph-2K using our previously published method [7, 43]. Complex I respiration was defined as malate/pyruvate/glutamate–driven oxygen consumption in the presence of ADP, whereas Complex II respiration was defined as succinate-driven oxygen consumption.

### Statistical Analyses

Data are presented as mean±S.D. Prism 9.0.0 software (GraphPad) was used for statistical analysis. To determine whether sample data has been drawn from a normally distributed population, D’Agostino–Pearson omnibus test or Shapiro–Wilk test was performed. For parametric data, ANOVA with post hoc Tukey comparison was used for multiple group comparison and t test was used to compare two different groups. For nonparametric data, Mann–Whitney U test was used. The threshold of P<0.05 was set to consider data statistically significant.

### Data availability

Raw data and complete MS data sets have been uploaded to the Center for Computational Mass Spectrometry, to the MassIVE repository at UCSD, and can be downloaded using the following link: https://massive.ucsd.edu/ProteoSAFe/dataset.jsp?task=6dfcd4ae92a7471a85fa0bf0add8c9bd (M assIVE ID number: MSV000091214; ProteomeXchange ID: PXD039904). [Note to the reviewers: To access the data repository MassIVE (UCSD) for MS data, please use: Username: MSV000091214_reviewer; Password: winter].

## Results

DCAs are activated to coenzyme A (CoA) by an as-yet-unknown acyl-CoA synthase enzyme and chain-shortened, preferentially within peroxisomes. In *ex vivo* studies [44, 45], dodecanedioc acid, a 12-carbon DCA (DC_12_), was chain-shortened as far as DC_4_-CoA, better known as succinyl-CoA [44, 45]. Because succinyl-CoA is a highly reactive metabolite that readily modifies lysine residues, we leveraged the succinyllysine PTM as a biomarker to establish which tissues take up and catabolize DCAs. First, we fed wild-type mice with diets containing either 10% w/w dodecanedioic acid (DC_12_), 10% w/w decanedioc acid (DC_10_), or 10% w/w octanedioic acid (DC_8_) mixed into the standard laboratory rodent diet. After 7 days consuming DCAs (Figure 1A), western blotting with a pan anti-succinyllysine antibody revealed that DC_10_ and DC_12_ induced a massive upregulation of this PTM in both liver and kidney, two tissues rich in peroxisomes (Figure 1B). Interestingly, DC_8_ induced protein succinylation only in kidney (Figure 1B), possibly because its shorter fatty acid chain stimulated other enzymatic pathways. Muscle and heart, which have few peroxisomes, showed no obvious change in protein succinylation by Western blot (data not shown). We chose DC_8_ and DC_12_ for further study. Consuming these DCAs did not affect cellular morphology of either liver or kidney, as evidenced by the H&E-stained tissues (Figure 1C). To test for the timing of hypersuccinylation and washout we performed a timed feeding experiment where food was provided and removed at regular intervals. Here we found that succinylation was maximal around 72 hours upon feeding. Following a three-day washout, succinylation levels were dramatically decreased. Moreover, a five-day washout was sufficient to return succinylation levels to near-baseline, suggesting that the kidney proteome is turned over during this timeframe (Figure 1D). This is key as it provides a therapeutic window that allows for widespread use of this AKI therapeutic, during the 72-hour window when injury is known to peak after AKI and likely leads to minimal side effects due to the fast washout time.

**Figure 1.**
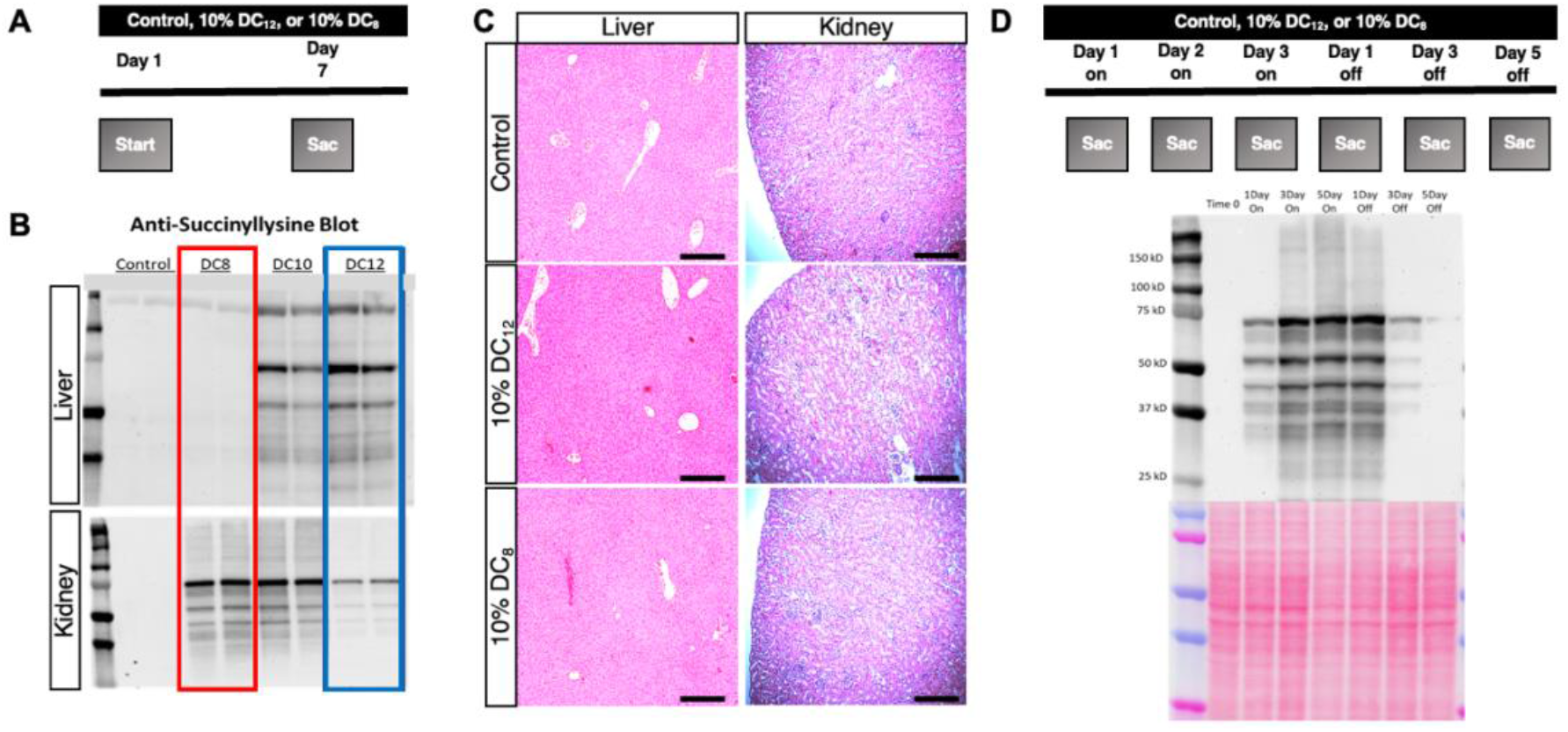
DC_8_ promotes succinylation in kidneys only whereas DC_12_ does it in both kidneys and liver. **A)** Schematic showing mouse supplementation regimen with a 10% DC_8_ w/w, 10% DC_10_ w/w, or 10% DC_12_ w/w for one week. **B)** Western blotting showing expression pattern of succinyllysine in the livers and kidneys of the supplemented animals; **C)** No differences observed in the liver or kidney morphology of control, 10% DC_12_-, and 10% DC_8_-fed animals; **D)** Five days upon termination of the medium-chain fatty acid-enriched diet, succinylation expression is vanished from the body, evidencing no risk on long-term accumulation. N=2. Scale bars represent a 5x magnification.

Next, given the known protection of peroxisomal FAO during AKI, we tested the DCA diets in the context of ischemia-reperfusion injury (IRI). Mice were fed with either control, 10% DC_12_, or 10% DC_8_-supplemented diets for seven days before and after IRI (Figure 2A). Upon tissue harvesting, control kidneys were much paler than the kidneys from DCA-fed groups, suggesting the latter group maintains a healthier appearance (Figure 2B). Moreover, DCA supplementation conferred an extensive upregulation of renal succinyllysine in both contralateral healthy and injured kidneys (Supplementary Figure 1). Our results further demonstrate that in comparison to control-fed mice, DCA-fed animals were significantly protected, as shown by significantly lower levels of serum creatinine and blood urea nitrogen (BUN) (Figure 2C). DCA-fed animals also presented improved renal morphology consistent with protection, including a marked decline in tubular dilation, tubular cast formation and a higher preservation of brush borders in comparison to control-fed animals (Figure 2D). More importantly, DC_8_-fed animals showed enhanced protection compared with DC_12_- fed mice (Figure 2B-D). The biochemical markers of AKI in mice supplemented with DC_8_ were very close to levels observed in sham-operated animals (Supplementary Figure 2A), and this was not caused by any changes in bodyweight (Supplementary Figure 2B). DC_8_-fed animals also presented fewer TUNEL positive cells (Figure 3A). In addition, the kidneys of DC_12_-fed animals had downregulation of the renal injury marker neutrophil gelatinase-associated lipocalin 2 (NGAL) and retained expression of a tubular cell marker (LTL). However, these changes were even more pronounced in DC_8_-fed animals (Figure 3B and 3B’), demonstrating that dicarboxylic acid supplementation prevents tubular cell damage, especially with the shorter length DCA, DC_8_. Given that DC_8_ was associated with enhanced protection from AKI, as well as the succinylation being confined to the kidney, we proceeded with DC_8_ in all remaining experiments.

**Figure 2.**
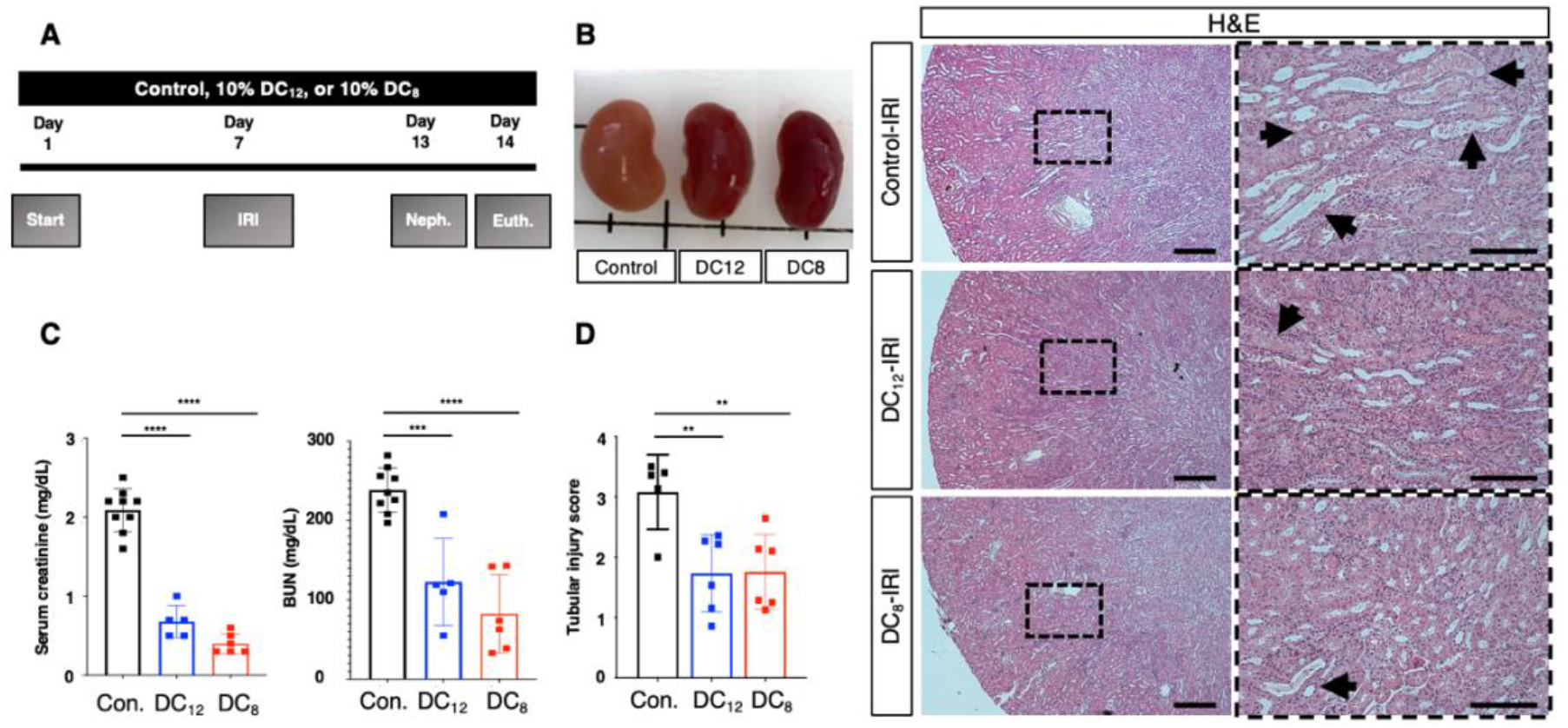
DC_12_ and DC_8_ supplementation protect male mice from ischemia-reperfusion induced AKI. **A)** Schematic showing mouse supplementation regimen with a 10% DC_8_ w/w or 10% DC_12_ w/w, as well as the IRI procedure and euthanasia; **B)** Anatomical kidney differences of control-, DC_12_-, DC_8_-supplemented animals upon tissue harvesting; **C)** Serum creatinine and BUN levels; **D)** H&E staining evidencing morphological changes in dicarboxylic-acid supplemented animals, such as a decline in tubular cast formation, dilation, and mitigated brush border loss, especially in DC_8_ animals. N=5-9. Scale bars represent a 10x and 20x magnification, respectively. Results are expressed as mean +- S.D. Prism 9.0.0 software (GraphPad) was used for statistical analysis. Analysis was performed using Student’s t test. Significance was given by a p value < 0.05. *p<0.05 **p<0.01 ***p<0.001 ****p<0.0001.

**Figure 3.**
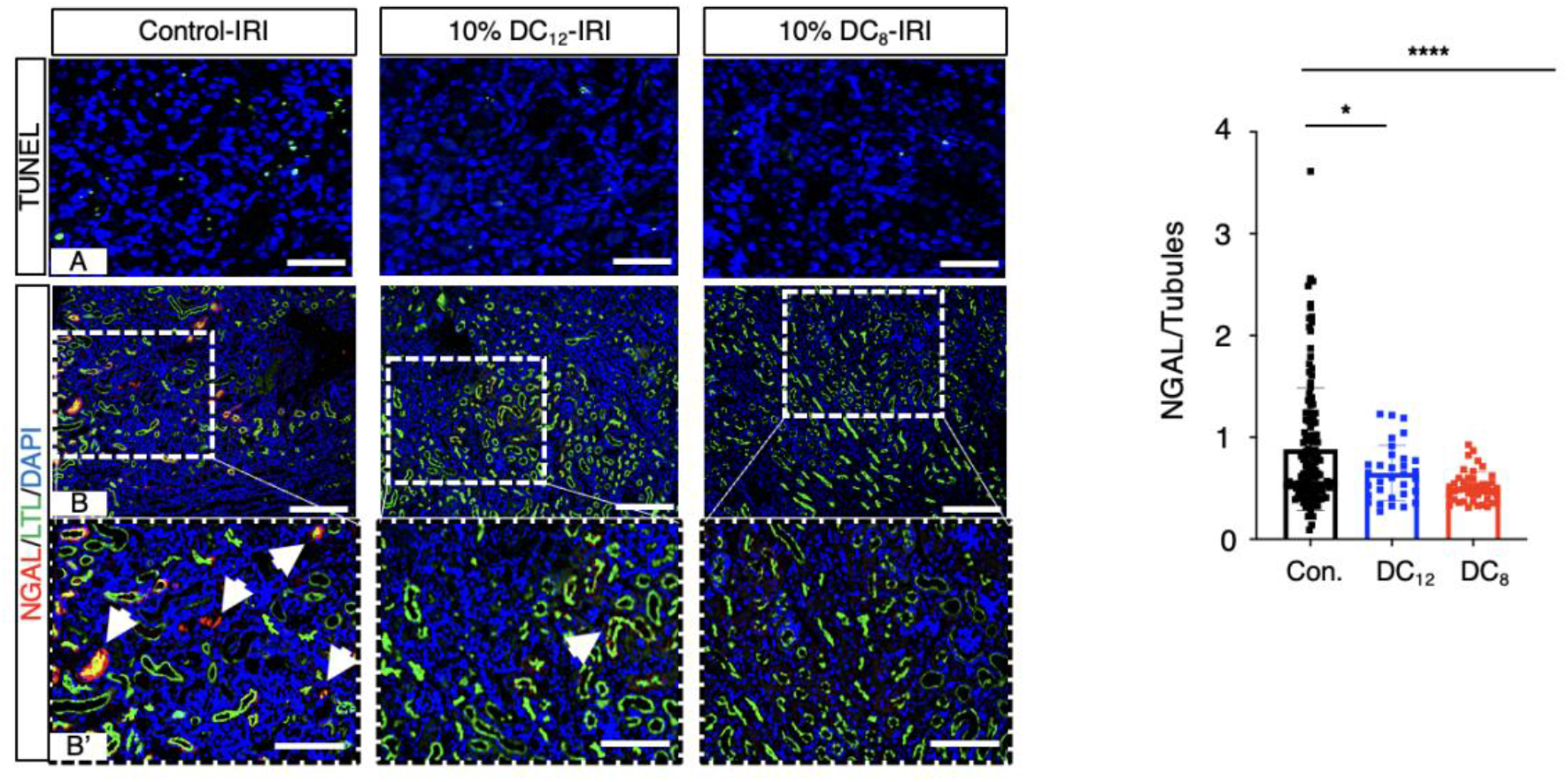
DC_12_ and DC_8_ supplementation mitigates tubular injury and prevents ischemia-reperfusion-induced AKI. **A)** TUNEL staining; **B)** renal expression of NGAL, LTL, and DAPI. N= 4. Scale bars represent a 10x magnification in images A-B and a zoomed area in B’. Results are expressed as mean +- S.D. Prism 9.0.0 software (GraphPad) was used for statistical analysis. Analysis was performed using Student’s t test. Significance was given by a p value < 0.05. *p<0.05 ****p<0.0001.

In another cohort of seven days-supplemented mice, an in-bolus cisplatin injection was performed, and the animals were sacrificed three days after (Figure 4A). In a similar protective manner, cisplatin-treated mice that were fed with DC_8_ were significantly protected from renal injury, as shown by lower levels of serum creatinine, reduced expression of renal NGAL at both the gene (Figure 4B, Supplementary Figure 3) and protein level (Figure 4D) and improved renal morphology, as shown by a decline in tubular dilation and cast formation, as well as minimized loss of brush borders (Figure 4C).

**Figure 4.**
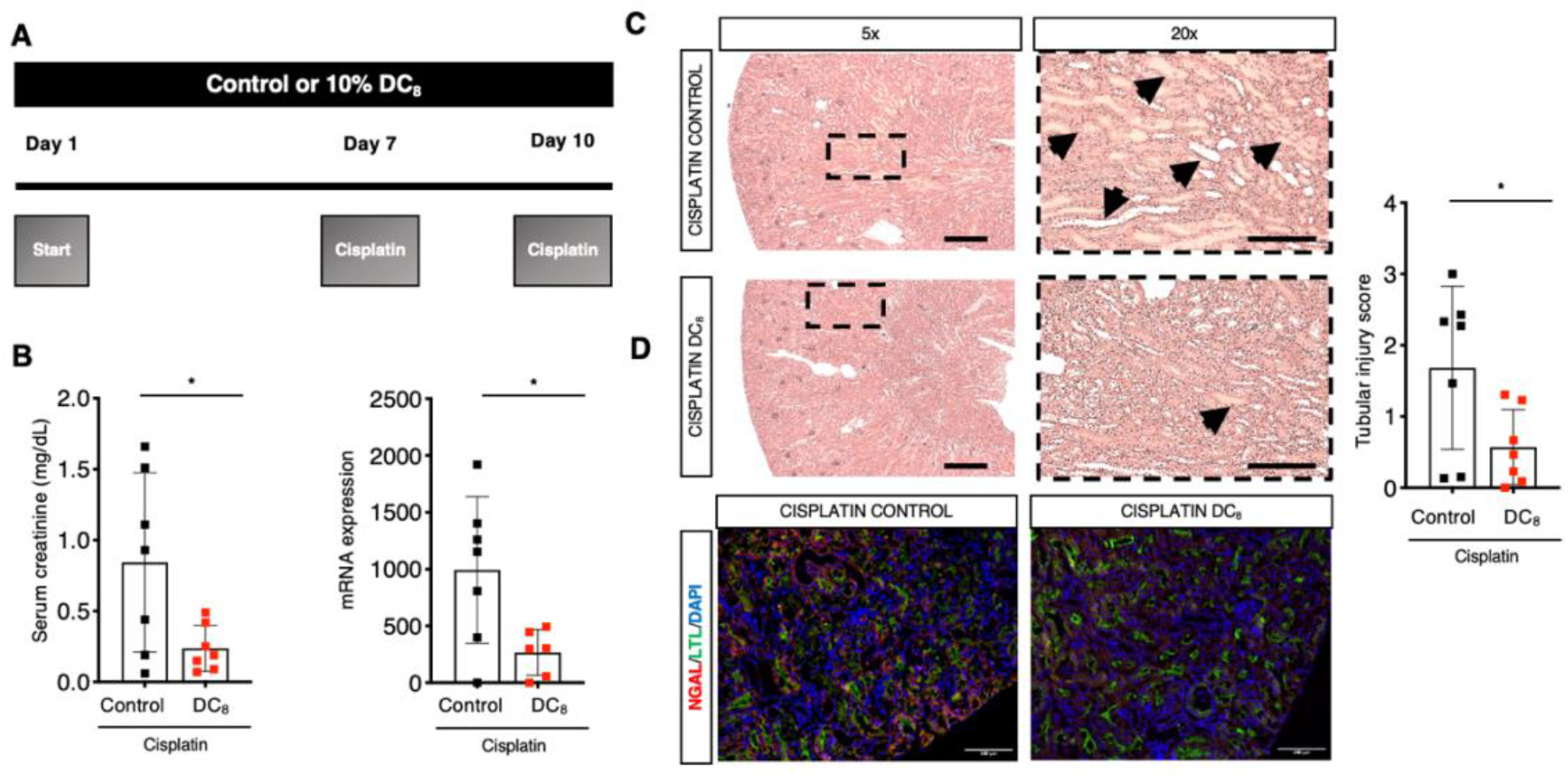
DC_8_-fed animals are protected from Cisplatin-induced AKI. **A)** Schematic of supplementation regimen, cisplatin injection, and euthanasia of animals; **B)** Serum creatinine and renal NGAL expression; **C)** H&E staining and tubular injury score (Scale bars represent a 5x and 20x magnification, respectively); **D)** immunofluorescence for NGAL, LTL, and DAPI. (Scale bars represent a 5x magnification). N=7 per group. Results are expressed as mean +- S.D. Prism 9.0.0 software (GraphPad) was used for statistical analysis. Analysis was performed using Student’s t test. Significance was given by a p value < 0.05. *p<0.05.

The initial choice of 10% w/w DCAs in the diet was informed by a previous study which compared the effects of 1.5% DC_10_ and 15% DC_10_ in a mouse model of type 2 diabetes [46]. We speculated that it would be possible to decrease this concentration without changing effectiveness. Therefore, we compared control-fed and animals supplemented with different concentrations of DC_8_, at 2.5%, and 5% w/w. Interestingly, while a dose of 2.5% w/w was protective compared to control-fed animals, it was suboptimal compared with 10%. However, a 5% DC_8_ supplementation was sufficient to protect mice from IRI-induced AKI to the same degree as a 10% supplementation (Figure 5A-B), further optimizing the translational potential in humans.

**Figure 5.**
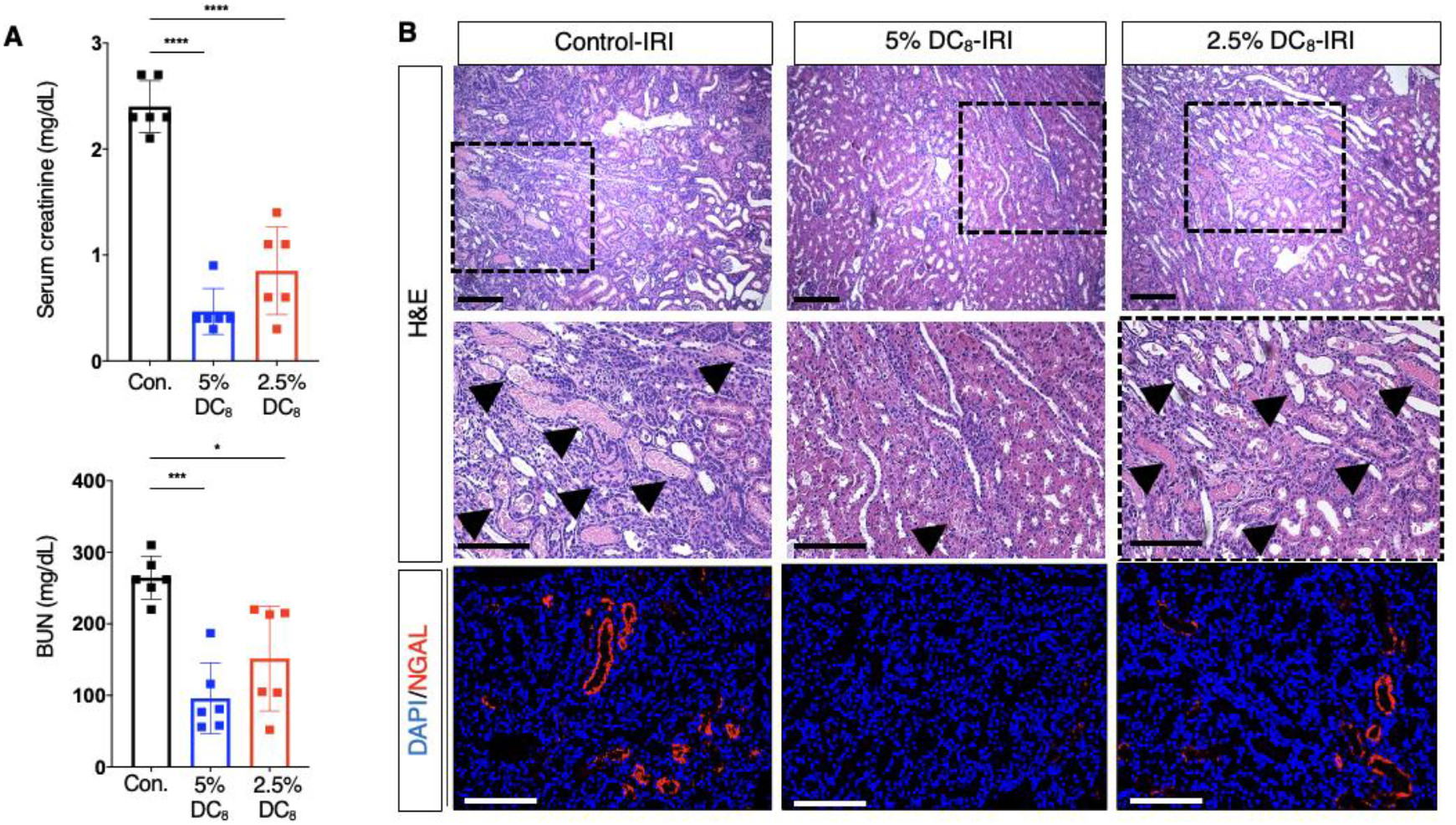
DC_8_ protects animals from AKI at a concentration as low as 5% w/w. **A)** Creatinine and BUN; **B)** H&E staining and immunofluorescence for NGAL and DAPI. N=6. Scale bars represent a 10x and 20x magnification, respectively. Results are expressed as mean +- S.D. Prism 9.0.0 software (GraphPad) was used for statistical analysis. Analysis was performed using Student’s t test. Significance was given by a p value < 0.05. *p<0.05 **p<0.01 ***p<0.001 ****p<0.0001.

Frozen injured and contralateral, uninjured kidneys of mice from the control and 10% DC_8_- supplemented groups (n = 4 per group) were analyzed via a comprehensive and sensitive label free lysine succinylation-based quantitative mass spectrometric workflow [28], that included an antibody-enabled succinylation enrichment, followed by data-independent acquisition (DIA) on an Orbitrap Eclipse mass spectrometer at the Buck Institute [25–27]. This workflow enabled the identification and quantification of 3,765 unique succinylation sites on 4,514 succinylated peptides from 1,160 succinylated protein groups (Supplementary Table 1), including 348 mitochondrial and 46 peroxisomal succinylated protein groups (Supplementary Figure 4A). The results highlight that DC_8_ dramatically remodeled the protein succinylome, a lysine PTM caused by interaction between proteins and the metabolite succinyl-CoA, formed by chain-shortening of DCAs (Figure 6A-D). The DC_8_ diet led to a drastic succinylation increase in the injured kidney with diet compared to control diet (Figure 6A), and a similarly strong succinylation increase was observed in the non-injured kidney (Figure 6B). Indeed, out of the 4,514 quantified succinylated peptides, 1,813 succinylated peptides (1,632 unique succinylation sites) were significantly altered with 1,686 up regulated succinylated peptides, when comparing DC_8_ vs control diet in injured kidneys (Figure 6A). With respect to non-injured kidneys, 1,606 succinylated peptides (1,471 unique PTM sites) were significantly changed with the DC_8_ diet, including 1,053 up regulated succinylated peptides (Figure 6B). More specifically, subcellular localization information revealed that DC_8_ induced a shift of the succinylation profiles of mitochondrial and peroxisomal proteins in injured kidneys, as increased succinylation levels were observed for 195 mitochondrial proteins and 44 peroxisomal proteins in the injured kidneys of DC_8_- vs control-fed mice (Supplementary Figures 4A and 4B, Supplementary Figure 5). In addition, gene ontology (GO) analysis of the changes in succinylated proteins revealed that mitochondrial cellular components were down-regulated in the injured kidneys of DC_8_-fed in comparison to control-fed animals (Figure 6C), while cellular components related to peroxisome and fatty acid metabolism were up-regulated (Figure 6D).

**Figure 6.**
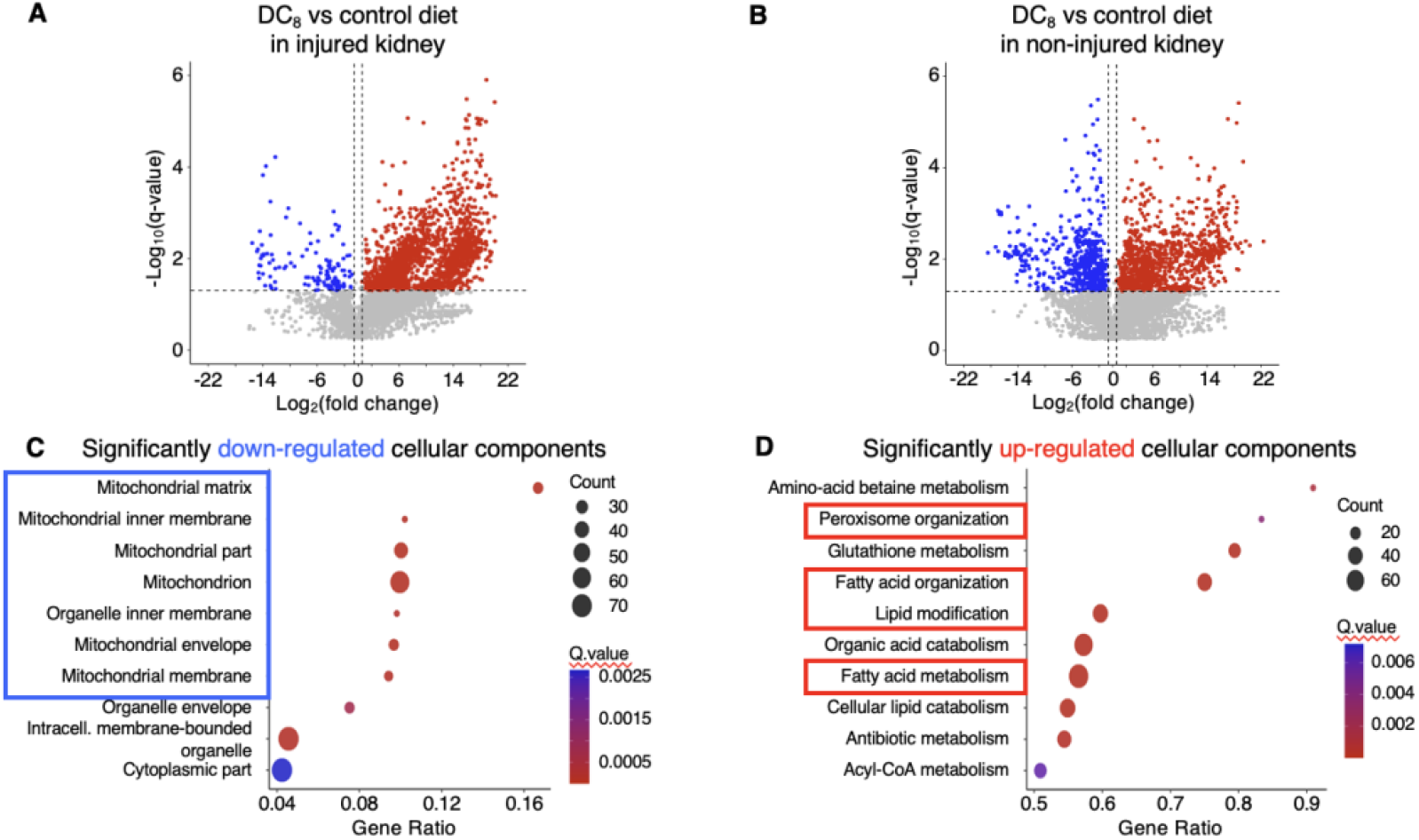
Lysine succinylome-based mass spectrometry evidences kidneys present a huge succinylome remodeling with DC_8_. **A)** Comparison between injured and **B)** non-injured kidneys of DC_8_ and control diet; **C)** downregulated and **D)** upregulated cellular components (top10 displayed) in injured kidneys from DC_8_-fed in comparison to control-fed animals. N= 4 per group.

Further data evidenced that increased peroxisomal succinylation is found in DC_8_-fed animals, even in baseline samples that were not subjected to IRI. Specifically, several key enzymes involved in the different steps of the peroxisomal FAO pathway were hypersuccinylated upon DC_8_ diet in the injured kidneys, as highlighted by both the increased ratios [displayed as log_2_ (DC_8_ vs control diet)] and the many succinylation sites observed per enzyme (Figure 7). Strikingly, the vast majority of the DC_8_-targeted lysine residues in mouse are also conserved in human, and for visualization those conserved sites are indicated in orange, which is highly relevant for potential clinical translational applications.

**Figure 7.**
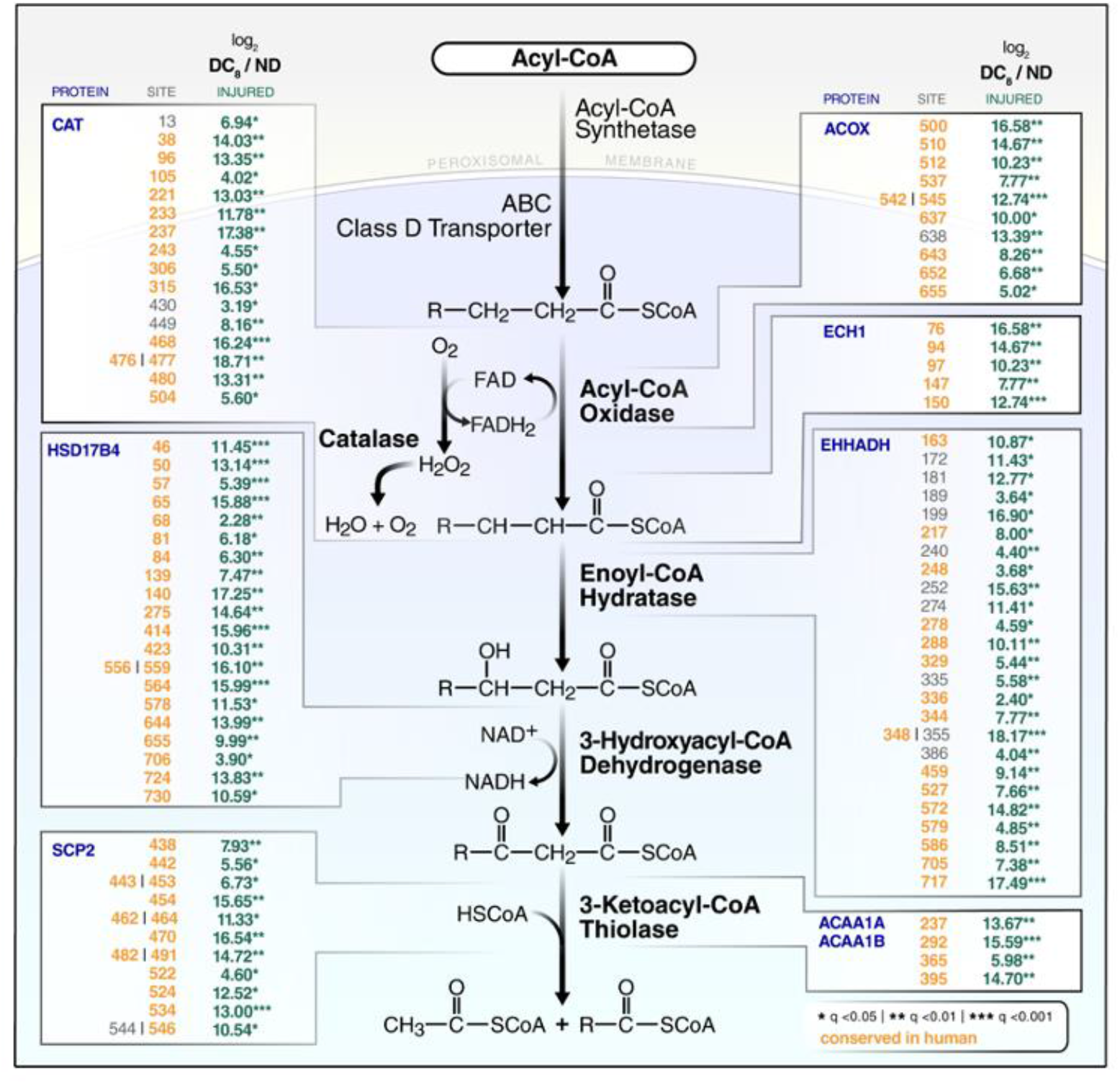
Peroxisomal enzymes and metabolites increased in the kidneys of DC_8_- fed animals. Several key enzymes involved in the different steps of the peroxisomal FAO pathway were hypersuccinylated upon DC_8_ diet in the injured kidneys ND: Normal/Control diet; DC_8_: animals fed with 10% DC_8_. N=4 per group.

With respect to uninjured, contralateral kidneys, proteomic analysis performed at the protein level (whole lysate analysis) showed that 89 proteins demonstrated a significant change in abundance (Q<0.01, FC>1.5) in uninjured, contralateral kidneys of DC_8_- vs control-fed mice (Supplementary Table 2). Of these, 66 were increased and 23 were decreased. Among the increased proteins were numerous peroxisomal enzymes required for DC_8_ catabolism, such as Abcd3, EHHADH, Hsd17b4, Acaa1a, Acaa1b, and Acot4. Reactome pathway analysis indicated a clear dichotomy between the increased and decreased proteins. Increased proteins in DC_8_ kidneys following AKI clustered in metabolic pathways in both mitochondria and peroxisomes (Supplementary Figure 6), while downregulated proteins were involved in cellular processes such as signaling (protein phosphorylation), extracellular matrix remodeling, and immune activation (Supplementary Figure 6).

Regardless of statistical significance of the whole protein lysate analysis, the abundance of all peroxisomal proteins quantified in DC_8_ against normal diet in non-injured (contralateral) kidneys showed that there was an overall trend toward increased peroxisomal protein abundance (∼1.4- fold on average) (Figure 8A), suggesting that DC_8_ increased the abundance of peroxisomes in order to promote its own catabolism. Functionally, we observed an increase in ^16C^-palmitate oxidation in kidneys from DC_8_-fed mice, particularly in the peroxisomal compartment (Supplementary Figure 7). Interestingly, two endoplasmic reticulum cytochrome proteins thought to be rate-limiting for the production of endogenous DCAs from medium-chain FAs, Cyp4a10 and Cyp4a14, were among the most deregulated proteins, with increases of 3.0-fold and 9.5-fold, respectively, in the contralateral kidneys of DC_8_- vs control-fed animals (Supplementary Table 2. There was also a minor shift upward in the abundance of the mitochondrial proteome (Fig 8B), but with only a few exceptions, these changes did not reach statistical significance.

**Figure 8.**
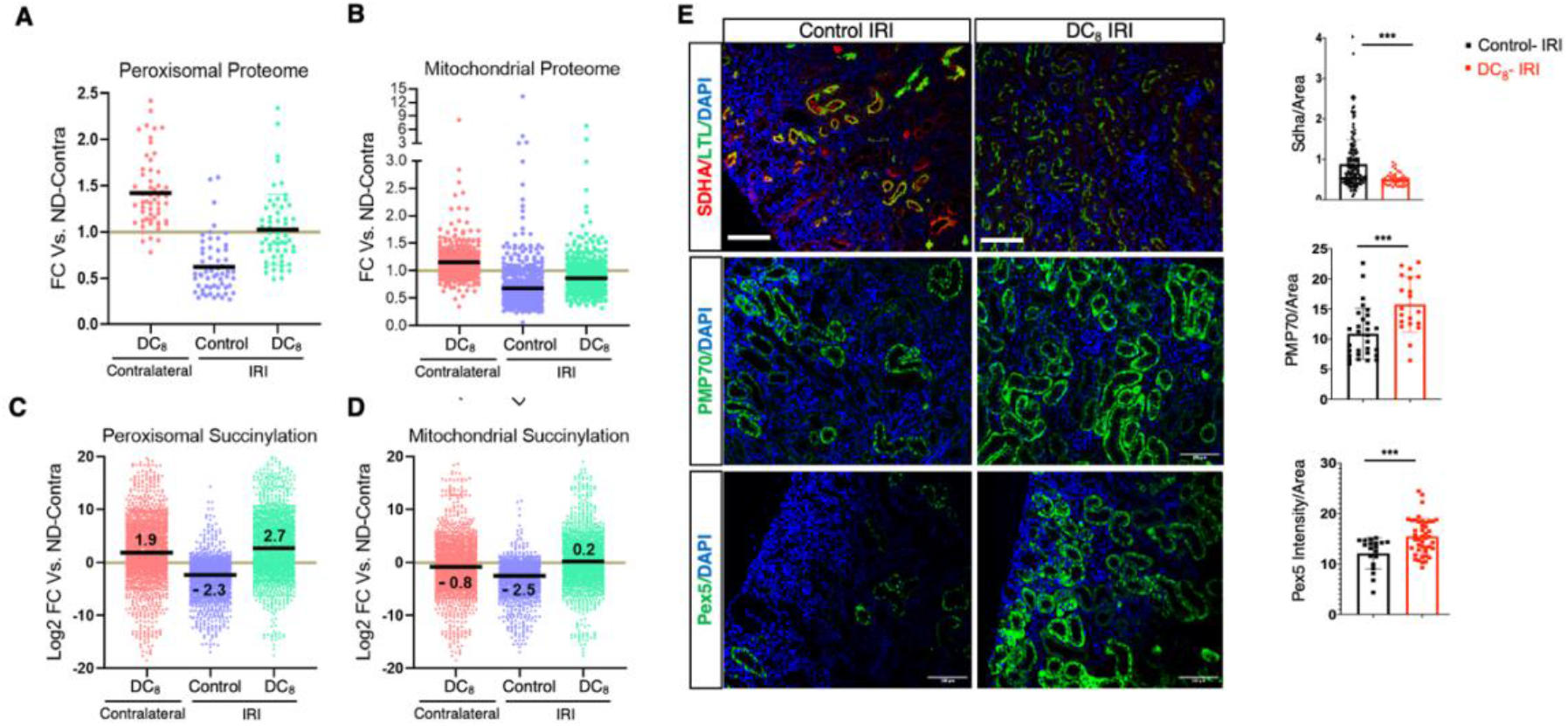
Supplementation with DC_8_ promotes peroxisomal succinylation, especially upon renal insult. Comparison of **A)** peroxisomal proteome, **B)** mitochondrial proteome, **C)** peroxisomal succinylation, and **D)** mitochondrial succinylation among non-injured kidneys fed with DC_8_ or control/normal diet (ND) and injured kidneys fed with DC_8_. **E)** immunofluorescence for the oxidative marker SDHA, and the peroxisomal markers PMP70 and PEX5. Scale bars represent a 5x and 20x magnification, respectively. N= 5. Results are expressed as mean +- S.D. Prism 9.0.0 software (GraphPad) was used for statistical analysis. Analysis was performed using Student’s t test. Significance was given by a p value < 0.05. *p<0.05 **p<0.01 ***p<0.001.

Succinylated peptide-level proteomic analysis identified 571 unique succinylation sites across 49 peroxisomal proteins (Supplementary Table 1). Feeding DC_8_ shifted the mean intensity of succinylation in the peroxisome upward about 4-fold (Log2FC 1.9) relative to normal diet uninjured kidneys (Fig 8C). At the same time, succinylation (and therefore succinyl-CoA) is modestly reduced in mitochondria from DC_8_-fed non-injured kidneys (Figure 8D). Succinylation is also shown to be increased in IRI-operated kidneys, and this is further confirmed by the markers peroxisomal biogenesis factor 5 (pex5) and peroxisomal membrane protein (pmp70), which were significantly increased in DC_8_-fed animals. Meanwhile, succinate dehydrogenase complex flavoprotein subunit A (Sdha), which codes one of the four subunits of the succinate dehydrogenase enzyme, was significantly decreased (Figure 8E). These data suggest that DC_8_ is predominantly oxidized in peroxisomes. The suppression of lysine succinylation in mitochondria may be caused by transfer of succinate from peroxisomes to mitochondria, which would enter the TCA cycle downstream of succinyl-CoA, thereby reducing endogenous succinyl-CoA levels and thus protein succinylation in the mitochondria.

At the earliest stages of this study, we hypothesized that feeding mice peroxisome-targeted fatty acids, in the form of DCAs, would promote the peroxisomal β-oxidation pathway. Indeed, we observed significantly higher peroxisomal oxidation of ^16^C-palmitate in non-injured (contralateral) DC_8_-fed kidney homogenates (Supplementary Figure 7). Proteomic analyses revealed that, in injured kidneys, AKI reduced the overall relative abundance of both peroxisomal and mitochondrial proteins (Fig 8A, B) (whole lysate analysis), as well as succinylation in both compartments (Fig 8C, D) (succinyl-enrichment analysis). These changes were largely normalized by the DC_8_ diet. With regards to peroxisomal protein succinylation, ‘DC_8_ + injury’ increased succinylation levels even higher than ‘DC_8_ + no-injury’, when compared to normal diet in contralateral kidneys, suggesting enhanced flux of DC_8_ through this pathway following AKI. Whereas DC_8_ diet alone significantly changed abundance of just 89 proteins, following injury, this increased to 706 proteins, of which 379 were significantly increased and 327 were decreased (Supplementary Table 2).

Functionally, we observed a nonsignificant trend toward higher C_16_-palmitate oxidation in injured kidneys from DC_8_-fed, whereas contralateral kidneys from DC_8_-fed were significantly different (Figure 9A-D). This assay relied upon etomoxir, an inhibitor of mitochondrial FAO, to discriminate between the two pathways. To further explore FAO, we performed ^14^C-flux assays with two peroxisome-specific substrates (^14^C^-^C_24_ and ^14^C-DC_12_) and a mitochondrial-specific substrate (^14^C-C_8_). For all three substrates, the rate of flux was significantly higher in injured kidneys from DC_8_-fed animals (Fig 9E-G).

**Figure 9.**
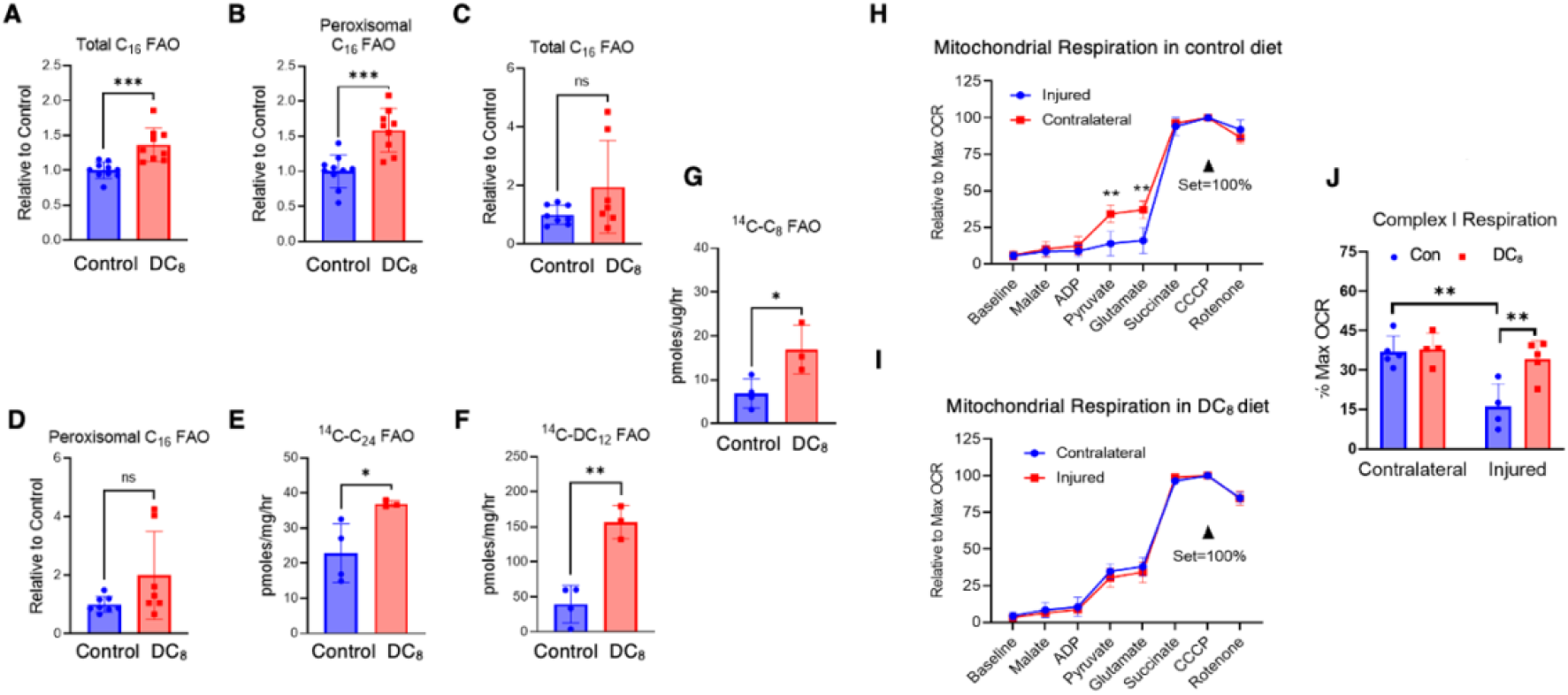
DC_8_ protects from AKI via increased peroxisomal FAO and changes in mitochondrial respiration. **A-G)** Comparison between the kidneys of control- and DC_8_-fed animals in terms of different peroxisomal FAO targets (A-B contralateral kidneys, C-G injured kidneys); **H)** Mitochondrial respiration in contralateral and injured kidney homogenates from control- and **I)** DC_8_-fed animals; **J)** complex I respiration among the groups. N= 5. Results are expressed as mean +- S.D. Prism 9.0.0 software (GraphPad) was used for statistical analysis. Analysis was performed using Student’s t test. Significance was given by a p value < 0.05. *p<0.05 **p<0.01 ***p<0.001.

Finally, we used oroboros high-resolution respirometry to measure respiratory-chain function in both contralateral and injured kidney tissue. For these assays, mitochondria were isolated and compared head-to-head, normalized first to protein concentration and then to the maximum, uncoupled rate of oxygen consumption to allow data from multiple sessions to be combined and compared. Thus, differences observed in these experiments are independent of any changes in mitochondrial number. In normal diet-fed mice, we observed a striking effect of AKI on the rate of oxygen consumption induced by the Complex I (NADH-producing) substrates pyruvate and glutamate (Figure 9H), meanwhile DC_8_-fed kidneys exhibited complete protection against this AKI induced loss of Complex I respiration (Figure 9I). Summarizing these data in bar graph format (Figure 9J), it can be seen that the DC_8_ diet has no effect on Complex I respiration in contralateral kidneys and is resistant to the loss of activity following AKI.

## Discussion

Dysregulated energy metabolism has been recognized as a key component of AKI for many years. Many attempts have been made at increasing mitochondrial function to limit injury, and none have translated to the clinic. Proximal tubules are rich in peroxisomes [47–49]. Over 40 years ago it was noted that AKI badly damages peroxisomes [50–52]. More recently, several studies indicated that bolstering peroxisomal FAO can protect against AKI. For example, inhibiting mitochondrial FAO with etomoxir leads to increased compensatory peroxisomal FAO and protection against AKI [18]. Similarly, sirtuin-1 (Sirt1) over-expression in mice protects against AKI by increasing peroxisome abundance and function [53]. Ablating sirtuin-5 (Sirt5) causes reduced mitochondrial FAO and compensatory peroxisomal FAO, with protection against AKI [7]. Finally, drugs that activate peroxisome proliferator activated receptor-alpha (PPARα), a transcription factor that drives peroxisomal biogenesis and FAO, are protective against AKI [18, 54–60]. Despite these promising studies, there has not been an effective non-genetic, non-toxic therapy to enhance peroxisomal FAO.

The proximal tubule is one of the most energy demanding cell types of the kidney, being very rich in both mitochondria and peroxisomes, and therefore the primary site of damage after ischemia [61, 62]. Peroxisomes are significantly damaged after a kidney injury, and unlike mitochondria, are much slower to regenerate. The loss of peroxisomes deprives the proximal tubules of their capacity to efficiently neutralize Reactive Oxygen Species (ROS) and any peroxides derived from normal metabolism. These findings are not limited to ischemic AKI: several studies confirm that the nephrotoxin cisplatin halts cellular energy production and accumulates in mitochondria, therefore affecting mostly proximal tubules at a rate five times higher than in bloodstream [63–65]. Nonetheless, cisplatin also suppresses peroxisome proliferator-activated receptor alpha (PPARα) activity, and a myriad of PPARα agonists have shown a protective effect when administered before cisplatin, demonstrating that promotion of peroxisomal activity is sufficient to protect from this drug-induced form of AKI [59, 60, 65]. Furthermore, sepsis promotes a decline in the peroxisomal population, culminating with a disfunction in its activity, especially in the import of peroxisomal targeting signal type 1 [66].

In this study we aimed to harness this protective effect of increased peroxisomal activity by supplementing these organelles with preferred substate of dicarboxylic acids. Increased peroxisomal activity uses much less oxygen per unit of energy and creates succinate, which is a simpler (and faster to metabolize) energy source for the mitochondria, all of which limit damaging ROS production. We postulate that due to these physiological and cellular circumstances, AKI has the most deleterious effect on the proximal tubules due to the loss of the peroxisomal activity. Mitochondria have long been the focus of renal-targeted therapies, but we show in this study that a profound protective effect can be elicited by promoting peroxisomal activity.

In the liver, DCAs were shown to be chain-shortened to succinyl-CoA inside the peroxisome, with subsequent hydrolysis by acyl-Coa Thioesterase 4 (ACOT4) and transfer of succinate to mitochondria [9–11]. Succinyl-CoA is traditionally thought of as a mitochondria-specific metabolite, an intermediate in the TCA cycle. Additionally, succinyl-CoA is known to non enzymatically acylate lysine residues of proteins, resulting in lysine succinylation [67, 68]. Succinyl CoA is very labile and difficult to measure [69]; we believe that lysine succinylation serves as a proxy for succinyl-CoA, with mitochondrial succinylation serving to indicate TCA cycle activity, and peroxisomal succinylation serving to indicate DCA catabolism.

Interestingly, although the succinate metabolized in peroxisomes is transferred to mitochondria, our study shows that DC_8_ does not undergo any hepatic metabolism, as its succinylation occurs exclusively in the kidneys. Given that this was the only physiological difference observed between DC_12_ and DC_8_, we speculate that the renal enzymes are crucial for the localized protection against AKI.

Additionally, we propose the protective effect is stimulated through the succinylation of key metabolic enzymes directly related to peroxisomally-derived succinate. Peroxisomally-derived succinate can be transported into the cytosol, unlike mitochondrial succinate. We have previously shown that deletion of the desuccinylase Sirt5 had a significantly protective effect after AKI [7]. After injury, Sirt5^-/-^ kidneys express hypersuccinylated mitochondrial enzymes, which decreases the mitochondrial FAO efficiency and enhances the effect of peroxisomal FAO. However, Sirt5 is a cytosolic protein and is not a readily druggable target. We now have found a more profound protective effect after AKI with DCA supplementation, which mechanistically does not decrease mitochondrial FAO, but rather enhances peroxisomal activity as dicarboxylic acids are direct substrates of peroxisomal FAO enzymes. The succinate produced by the peroxisomal FAO is transported to the mitochondrial as a source of fast energy, however, unlike succinyl-CoA produced by the mitochondria, this succinate is not enzymatically active. We suggest that DCA supplementation increasing peroxisomal activity to produce small direct substrates of mitochondrial FAO without otherwise damaging mitochondria (like in the Sirt5 knockouts) is a more cellularly efficient mechanism of protection. Additionally, diet supplementation rather than genetic knockout, is a more clinically translatable therapy.

In short, the concept of metabolically bolstering the kidney against injury with a naturally occurring lipid species is highly innovative and with a high degree of efficacy. We predict that DCAs will have wide utility for many forms of AKI and expect that DCA-induced metabolic protection will extrapolate to other organ injuries as well, thereby giving our studies wide scientific and therapeutic impact.

## Acknowledgments

This work is supported by NIH R01DK125015, NIH R01DK121758 (S.S.L), NIH R01DK090242 (S.S.L), NIH R21HD097403 (E.S.G), Richard K. Mellon Postdoctoral fellowship (K.P. and T.C), UPMC VMI Pilot Project Program in Hemostasis and Vascular Biology (T.C). We acknowledge the support of instrumentation for the Orbitrap Eclipse Tribrid system from the NCRR shared instrumentation grant 1S10 OD028654 (B.S.).

A.B., K.P., T.C., designed and performed experiments, provided intellectual input, and wrote the manuscript. A.B., K.P., T.C., J.B, J.P.R., J.B.B., C.D.K., A.O’B.R., U., V.Y., B. S. B. Z., B.S., A. V.S., E.S.G. performed experiments and provided intellectual input. B.S. and J.B. coordinated sample preparation with J.P.R. for mass spectrometric analysis followed by data acquisition by J.B. and C.D.K. J.B., J.B.B., A. O’B., and B.S. processed and interpreted mass spectrometry data and generated data results and graphical displays. E.S.G and S.S.L designed and performed experiments, edited the manuscript, provided intellectual input, and oversaw the project. All authors approved the final version of this manuscript.

We thank the UPMC Children’s Hospital of Pittsburgh Histology Core Laboratory and the Kansas State University Veterinary Diagnostic Laboratories for technical assistance.

## SUPPLEMENTARY APPENDIX

### Supplemental Methods

#### Proteomics Analysis

##### Protein digestion and desalting: Sample cohort

Both 7-day renal ischemia-reperfusion injury (IRI) and contralateral kidney tissues from control (n=4), 10% DC12 (n=3), and 10% DC8-fed (n=4) mice were collected. Frozen livers were immersed in 750 μL of lysis buffer containing 8 M urea, 200 mM triethylammonium bicarbonate (TEAB), pH 8, 75 mM sodium chloride, 1 μM trichostatin A, 3 mM nicotinamide, and 1x protease/phosphatase inhibitor cocktail (Thermo Fisher Scientific, Waltham, MA), and homogenized for 2 cycles with a Bead Beater TissueLyser II (Qiagen, Germantown, MD) at 24 Hz for 3 min each. Lysates were clarified by spinning at 15,700 x *g* for 15 min at 4°C, and the supernatant containing the soluble proteins was collected. Protein concentrations were determined using a Bicinchoninic Acid Protein (BCA) Assay (Thermo Fisher Scientific), and subsequently 5.1 mg of protein from each sample were aliquoted and samples were brought to an equal volume using a solution of 8 M urea in 50 mM TEAB, pH 8. Proteins were reduced using 20 mM dithiothreitol (DTT) in 50 mM TEAB for 30 min at 37 °C, and after cooling to room temperature, alkylated with 40 mM iodoacetamide (IAA) for 30 min at room temperature in the dark. Samples were diluted 4-fold with 50 mM TEAB, pH 8, and proteins were digested overnight with a solution of sequencing-grade trypsin (Promega, San Luis Obispo, CA) in 50 mM TEAB at a 1:50 (wt:wt) enzyme:protein ratio at 37°C. This reaction was quenched with 1% formic acid (FA) and the sample was clarified by centrifugation at 2,000 x g for 10 min at room temperature. Clarified peptide samples were desalted with Oasis 30-mg Sorbent Cartridges (Waters, Milford, MA). 100 μg of each peptide elution were aliquoted for analysis of protein-level changes, after which all desalted samples were vacuum dried. The 100 µg whole lysate aliquots were re-suspended in 0.2% FA in water at a final concentration of 1 µg/µL and stored for MS analysis. The remaining 5-mg digests were re-suspended in 1.4 mL of immunoaffinity purification (IAP) buffer (Cell Signaling Technology, Danvers, MA) containing 50 mM 4-morpholinepropanesulfonic acid (MOPS)/sodium hydroxide, pH 7.2, 10 mM disodium phosphate, and 50 mM sodium chloride for PTM enrichment. Peptides were enriched for succinylation with anti-succinyl antibody conjugated to agarose beads from the Succinyl-Lysine Motif Kit (Cell Signaling Technology). This process was performed according to the manufacturer protocol, however each sample was incubated in half the recommended volume of washed beads. Peptides were eluted from the antibody-bead conjugates with 0.15% trifluoroacetic acid (TFA) in water and were desalted using C_18_ stagetips made in-house. Samples were vacuum dried and re suspended in 0.2% FA in water. Finally, indexed retention time standard peptides (iRT; Biognosys, Schlieren, Switzerland) [29] were spiked in the samples according to manufacturer’s instructions.

##### Protein digestion and desalting: Spectral library of the mouse kidney whole lysate

To build a deep spectral library of the mouse kidney proteome (protein level), 1-mg of protein lysates of four different samples – one IRI lysate from a 10% DC12-fed mouse, one contralateral lysate from a 10% DC12-fed mouse, one from IRI lysate from a 10% DC8-fed mouse, and one contralateral lysate from a 10% DC8-fed mouse – was pooled to constitute a representative matrix. The sample pool was digested in-solution with trypsin and desalted with Oasis 30-mg Sorbent Cartridges as described above. The spectral library was generated by combining strong cation exchange (SCX) peptide fractionation, high pH reversed-phase (HPRP) peptide fractionation, and repetitive measurements of the unfractionated representative matrix.

For SCX peptide fractionation, 1 mg of desalted peptide elution was vacuum dried and resuspended in 25 mM sodium acetate, pH 5.5. Peptides were fractionated using Pierce Strong Cation Exchange Spin Columns (Thermo Fisher Scientific) according to manufacturer’s instructions, and 10 fractions were collected by sequentially adding 20 mM, 40 mM, 60 mM, 80 mM, 100 mM, 150 mM, 250 mM, 500 mM, 1000 mM, and 2000 mM NaCl in 25 mM sodium acetate, pH 5.5. Each elution fraction was desalted with Oasis 10-mg Sorbent Cartridges, samples were vacuum dried and re-suspended in 100 µL of 0.2% FA in water.

For the HPRP peptide fractionation, 100 µg of desalted peptide elution was vacuum dried and resuspended in 0.1% TFA. Peptides were fractionated using Pierce High pH Reversed-Phase Peptide Fractionation Kit (Thermo Fisher Scientific) according to manufacturer’s instructions, and eight fractions were collected by sequentially adding 5%, 7.5%, 10%, 12.5%, 15%, 17.5%, 20%, and 50% ACN in 0.1% triethylamine. Samples were vacuum dried and re-suspended in 12.5 µL of 0.2% FA in water.

For the unfractionated pool, the desalted peptide elution was vacuum dried and resuspended in 0.2% FA in water at a final concentration of 1 µg/µL.

Finally, iRT peptides [29] were spiked into all samples according to manufacturer’s instructions.

##### Mass Spectrometric Analysis: Sample Cohort

For the protein lysates (protein level analysis), LC-MS/MS analyses were performed on a Dionex UltiMate 3000 system online coupled to an Orbitrap Exploris 480 mass spectrometer (Thermo Fisher Scientific, San Jose, CA). The solvent system consisted of 2% ACN, 0.1% FA in water (solvent A) and 80% ACN, 0.1% FA in water (solvent B). Digested peptides (400 ng) were loaded onto an Acclaim PepMap 100 C_18_ trap column (0.1 x 20 mm, 5 µm particle size; Thermo Fisher Scientific) over 5 min at 5 µL/min with 100% solvent A. Peptides were eluted on an Acclaim PepMap 100 C_18_ analytical column (75 µm x 50 cm, 3 µm particle size; Thermo Fisher Scientific) at 300 nL/min using the following gradient of solvent B: linear from 2.5% to 24.5% in 125 min, linear from 24.5% to 39.2% in 40 min, up to 98% in 1 min, and back to 2.5% in 1 min. The column was re equilibrated for 30 min with 2.5% of solvent B, and the total gradient length was 210 min. Each sample was acquired in data-independent acquisition (DIA) mode. Full MS spectra were collected at 120,000 resolution (AGC target: 3e6 ions, maximum injection time: 60 ms, 350-1,650 m/z), and MS2 spectra at 30,000 resolution (AGC target: 3e6 ions, maximum injection time: Auto, NCE: 30, fixed first mass 200 m/z). The DIA precursor ion isolation scheme consisted of 26 variable windows covering the 350-1,650 m/z mass range with an overlap of 1 m/z (Supplementary Table 3) [27].

For the enriched succinylated peptides (PTM level analysis), LC-MS/MS analyses were performed on a Dionex UltiMate 3000 system online coupled to an Orbitrap Eclipse Tribrid mass spectrometer (Thermo Fisher Scientific). The solvent system consisted of 2% ACN, 0.1% FA in water (solvent A) and 98% ACN, 0.1% FA in water (solvent B). Succinylated-enriched peptides (4 µL) were loaded onto an Acclaim PepMap 100 C_18_ trap column (0.1 x 20 mm, 5 µm particle size; Thermo Fisher Scientific) over 5 min at 5 µL/min with 100% solvent A. Peptides were eluted on to an Acclaim PepMap 100 C_18_ analytical column (75 µm x 50 cm, 3 µm particle size; Thermo Fisher Scientific) at 300 nL/min using the following gradient of solvent B: 2% for 5 min, linear from 2% to 20% in 125 min, linear from 20% to 32% in 40 min, up to 80% in 1 min, 80% for 9 min, and down to 2% in 1 min. The column was equilibrated with 2% of solvent B for 29 min, with a total gradient length of 210 min. All samples were acquired in DIA mode. Full MS spectra were collected at 120,000 resolution (AGC target: 3e6 ions, maximum injection time: 60 ms, 350-1,650 m/z), and MS2 spectra at 30,000 resolution (AGC target: 3e6 ions, maximum injection time: Auto, NCE: 27, fixed first mass 200 m/z). The same isolation scheme as above was used (Supplementary Table 3) [27].

##### Mass Spectrometric Analysis: Spectral Library

For the 10 SCX fractions and the triplicate measurements of the unfractionated pool, LC-MS/MS analyses were performed on a Dionex UltiMate 3000 system online coupled to an Orbitrap Exploris 480 mass spectrometer operated in DIA mode, as described above, except that estimated 800 ng peptides were injected for the SCX fractions and 400 ng peptides for the unfractionated pool.

For the eight HPRP fractions, LC-MS/MS analyses were performed on a Dionex UltiMate 3000 system online coupled to an Orbitrap Eclipse Tribrid mass spectrometer operated in DIA mode, as described above, and 400 ng peptides were injected.

#### Data Analysis

##### Spectral library generation

A deep DIA spectral library of mouse kidney whole lysate was generated in Spectronaut (version 15.1.210713.50606; Biognosys) using BGS settings and a *Mus musculus* UniProtKB-TrEMBL database (86,521 entries, accessed on 08/24/2021). Briefly, for the Pulsar search, trypsin/P was set as the digestion enzyme and 2 missed cleavages were allowed. Cysteine carbamidomethylation was set as fixed modification, and methionine oxidation and protein N-terminus acetylation as variable modifications. Identifications were validation using 1% false discovery rate (FDR) at the peptide spectrum match (PSM), peptide and protein levels, and finally the best 3-6 fragments per peptide were kept. Finally, the library contains 55,552 modified peptides and 6,152 protein groups (Supplementary Table 4).

#### DIA Data Processing and Statistical Analysis

For the whole lysate (protein level) analysis, DIA data was processed in Spectronaut (version 15.1.210713.50606) using the previously described library. Data extraction parameters were set as dynamic and non-linear iRT calibration with precision iRT was selected. Identification was performed using 1% precursor and protein q-value. Quantification was based on the area of MS2 extracted ion chromatograms (XICs), and local normalization was applied. iRT profiling was selected.

For the PTM-enriched samples, DIA data was processing with directDIA embedded in Spectronaut (version 15.1.210713.50606) using the same mouse protein database as for generating the spectral library. Trypsin/P was set as digestion enzyme and two missed cleavages were allowed. Cysteine carbamidomethylation was set as fixed modification, and methionine oxidation, protein N-terminus acetylation and lysine succinylation as variable modifications. Data extraction parameters were set as dynamic. Identification was performed using 1% precursor and protein q-value. PTM localization was selected with a probability cutoff of 0.75. Quantification was based on the XICs of 3 – 6 MS2 fragment ions, specifically b- and y-ions, without normalization as well as data filtering using q-value sparse. Grouping and quantitation of PTM peptides were accomplished using the following criteria: minor grouping by modified sequence and minor group quantity by mean precursor quantity.

Differential expression analysis was performed using a paired t-test, and p-values were corrected for multiple testing, specifically applying group-wise testing corrections using the Storey method [30, 31]. For whole lysate analysis, protein groups are required with at least two unique peptides. For determining differential protein changes, a q-value < 0.01 and absolute Log_2_(fold-change) > 0.58 are required to qualify as ‘significantly-altered’ (Supplementary Table 2). For the PTM analysis, each succinylated peptide is quantified individually comparing conditions with significance cutoffs of q-value < 0.05, and absolute Log_2_(fold-change) > 0.58 (Supplementary Table 1).

#### Enrichment Analysis

An over-representation analysis was performed using ConsensusPathDB-mouse (Release MM11, 14.10.2021) [32, 33] to determine which gene ontology (GO) terms were significantly enriched. GO terms identified from the over-representation analysis were subjected to the following filters: q value < 0.01, count ≥ 3 and term level ≥ 3. Dot plots were generated using the ggplot2 package [34] in R.

##### Subcellular Localization

Subcellular localization was determined using Cytoscape [35] (version 3.8.2) and stringAPP [36] (version 1.7.0), by applying default settings, except that species was defined as *M. musculus*. Ten proteins for which no organelle information could be retrieved were excluded for the analysis, and only compartments with scores above 4.5 were considered.

##### LOPIT Organellar Protein Distribution

The hyperLOPIT2015 pluripotent stem cell dataset was downloaded in Rstudio (version 4.0.5, https://www.rstudio.com/ products/rstudio/download/#download, accessed on 31 March 2021) using the pRolocData R package (Bioconductor) and plotted with the pRoloc R package [37, 38].

To create the LOPIT organellar protein distribution dataset a crude membrane preparation from mouse pluripotent stem cell lysate was fractionated by density gradient ultracentrifugation to separate and enrich organelles based on density [37, 39, 40]. Proteins were identified via quantitative multiplexed mass spectrometry and assigned to organelles based on similarities in distribution in the density gradient to well-annotated organelle protein markers. The t-SNE machine learning algorithm was used to reduce the number of dimensions in the LOPIT proteomic data to a 2D map where proteins cluster by similarity from multiple experimental factors [41, 42]. All LOPIT protein assignments were used as originally determined and without additional refinement (Supplementary Figure 5).

### Supplementary Figures Legends

**Supplementary Figure 1:**
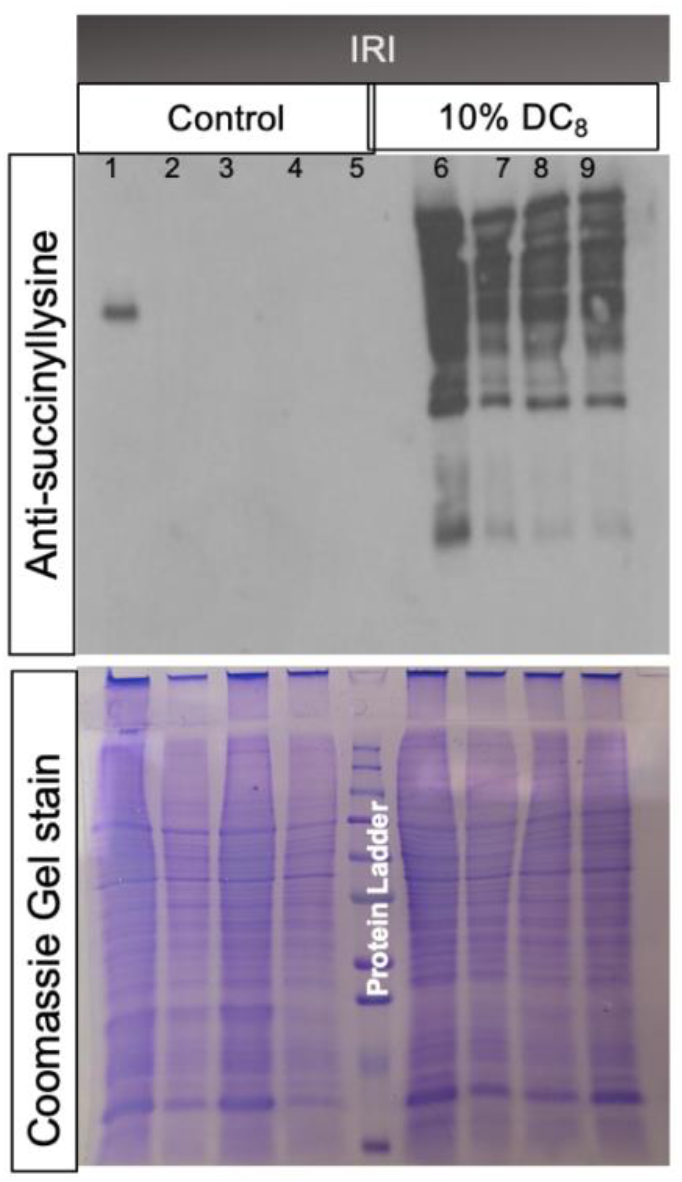
Kidney succinylation is increased in DC_8_-fed animals, both in contralateral healthy and in kidneys that were subjected to IRI. N=4.

**Supplementary Figure 2:**
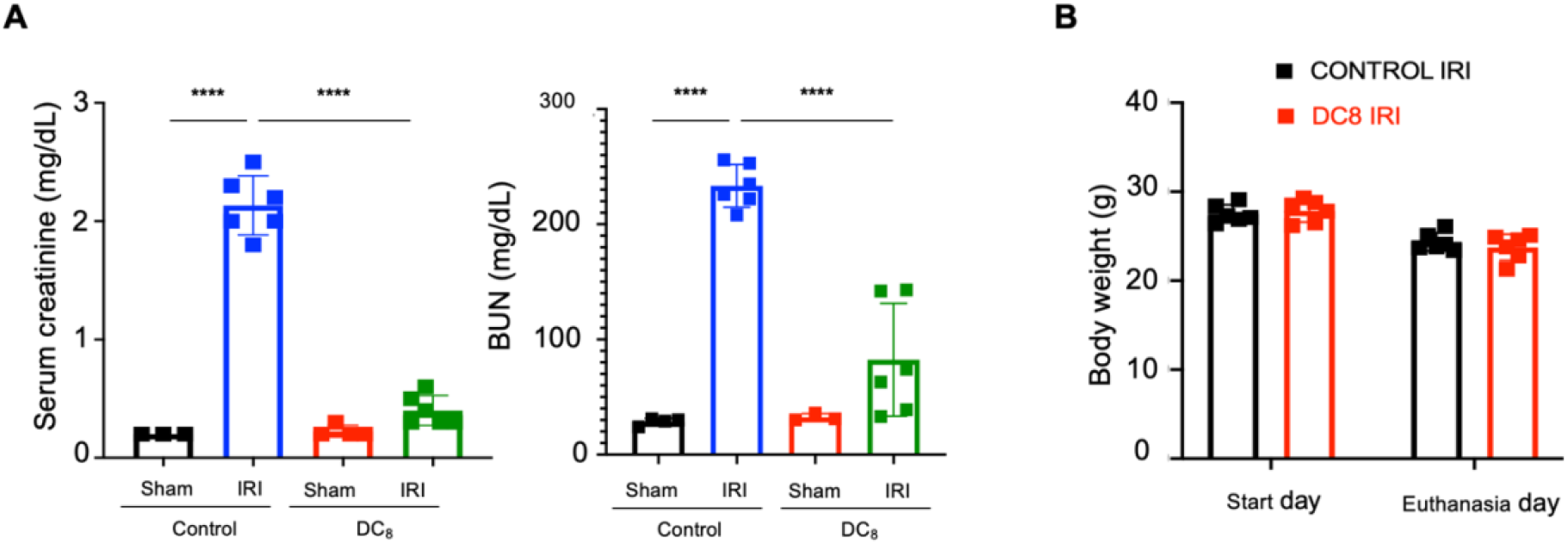
Comparison between Sham and IRI animals fed with either a control or DC_8_-diet. Creatinine and BUN show no significant differences between control- and DC_8_-fed animals that were solely subjected to a unilateral nephrectomy 24 hours prior to sacrifice, whereas when mice undergo IRI, DC_8_-fed are significantly protected from the injury. N= 6 (IRI) and 3 (sham). Results are expressed as mean +- S.D. Prism 9.0.0 software (GraphPad) was used for statistical analysis. Analysis was performed using Student’s t test. Significance was given by a p value < 0.05. *p<0.05 ****p< 0.0001.

**Supplementary Figure 3:**
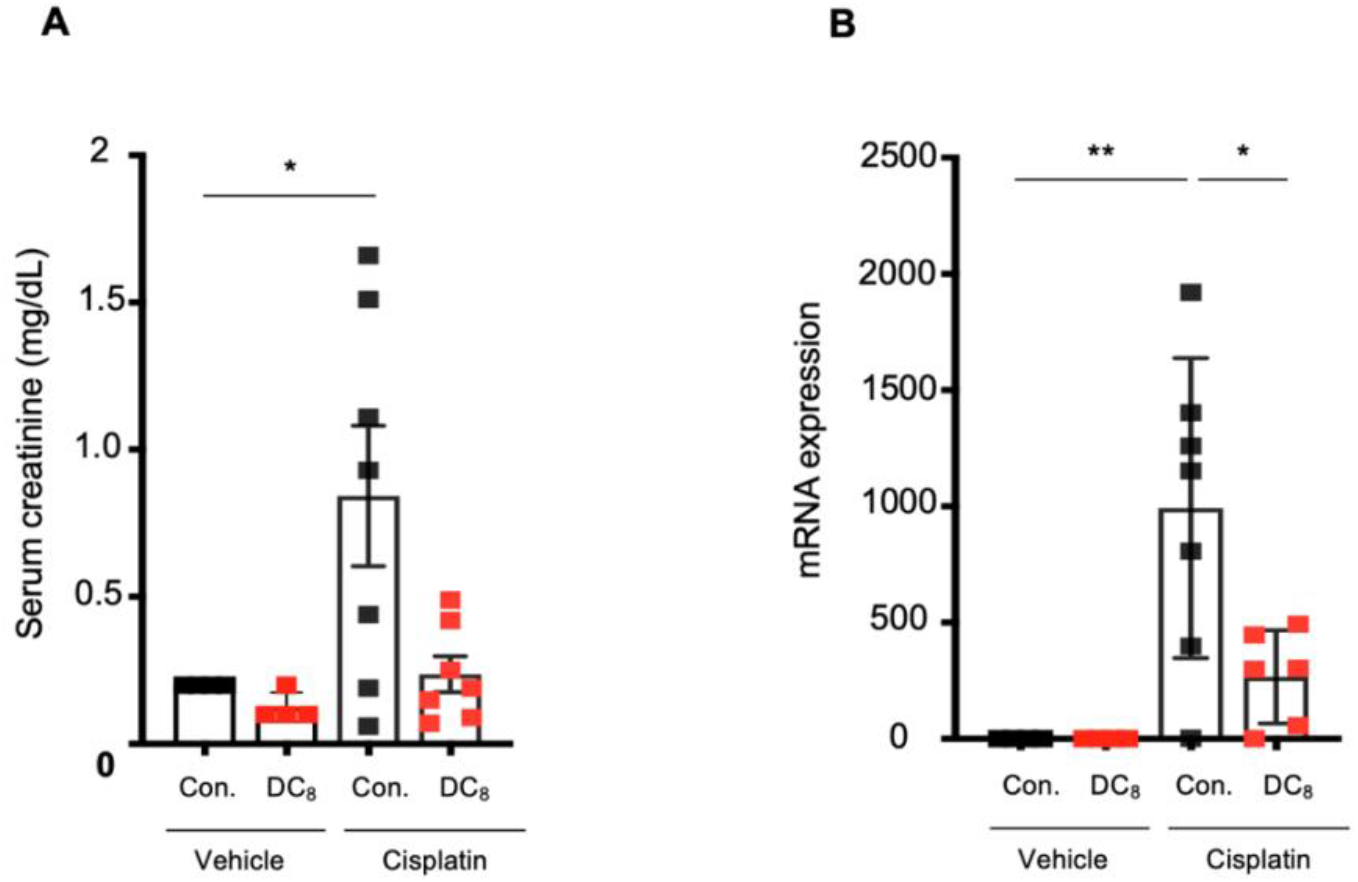
DC_8_-fed animals are protected from Cisplatin-induced AKI. A) Serum creatinine and renal NGAL expression; B) H&E staining. N=7 (cisplatin groups) and 4 (vehicle groups). Results are expressed as mean +- S.D. Prism 9.0.0 software (GraphPad) was used for statistical analysis. Analysis was performed using Student’s t test. Significance was given by a p value < 0.05. *p<0.05.

**Supplementary Figure 4:**
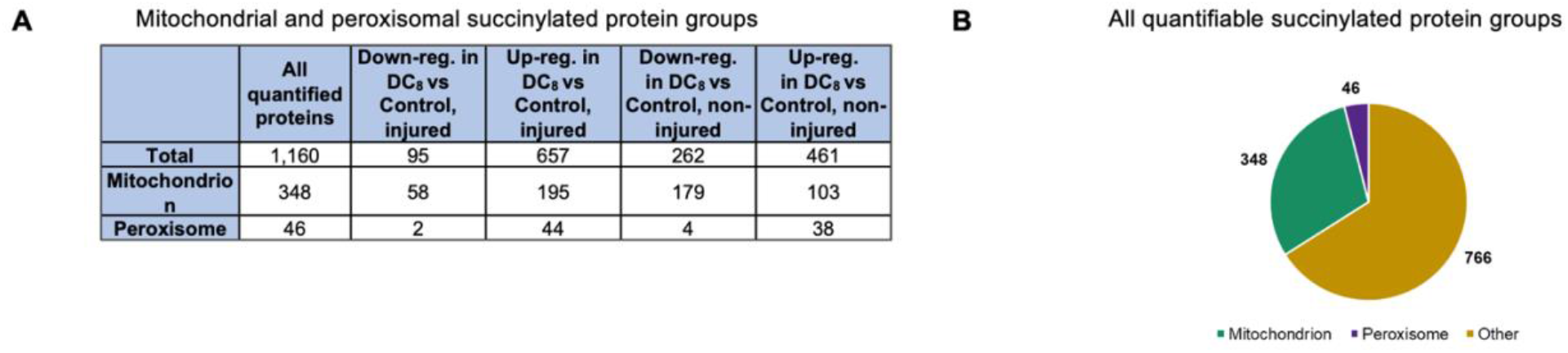
Lysine succinylome-based mass spectrometry evidences that DC_8_ induces a drastic remodeling of mitochondrial and peroxisomal succinylomes in both injured and non-injured kidneys. Protein subcellular localization information was retrieved using stringAPP (Szklarczyk *et al.*, 2015, *Nucleic Acids Res*, 43, D447-52), and the compartment score threshold was set at 4.5. ‘Total’ in **A** and ‘Other’ in **B** include proteins not assigned to mitochondrial nor peroxisomal compartments as well as proteins for which no subcellular localization information was retrieved. **A)** Summary of all succinylated proteins as well as mitochondrial and peroxisomal succinylated proteins that were quantified and significantly down- and up-regulated for DC_8_ vs Control comparisons in both injured and non-injured kidneys. **B)** Proportion of proteins assigned to mitochondrion (green), peroxisome (purple) and other organelles (beige) among all quantified succinylated protein groups.

**Supplementary Figure 5.**
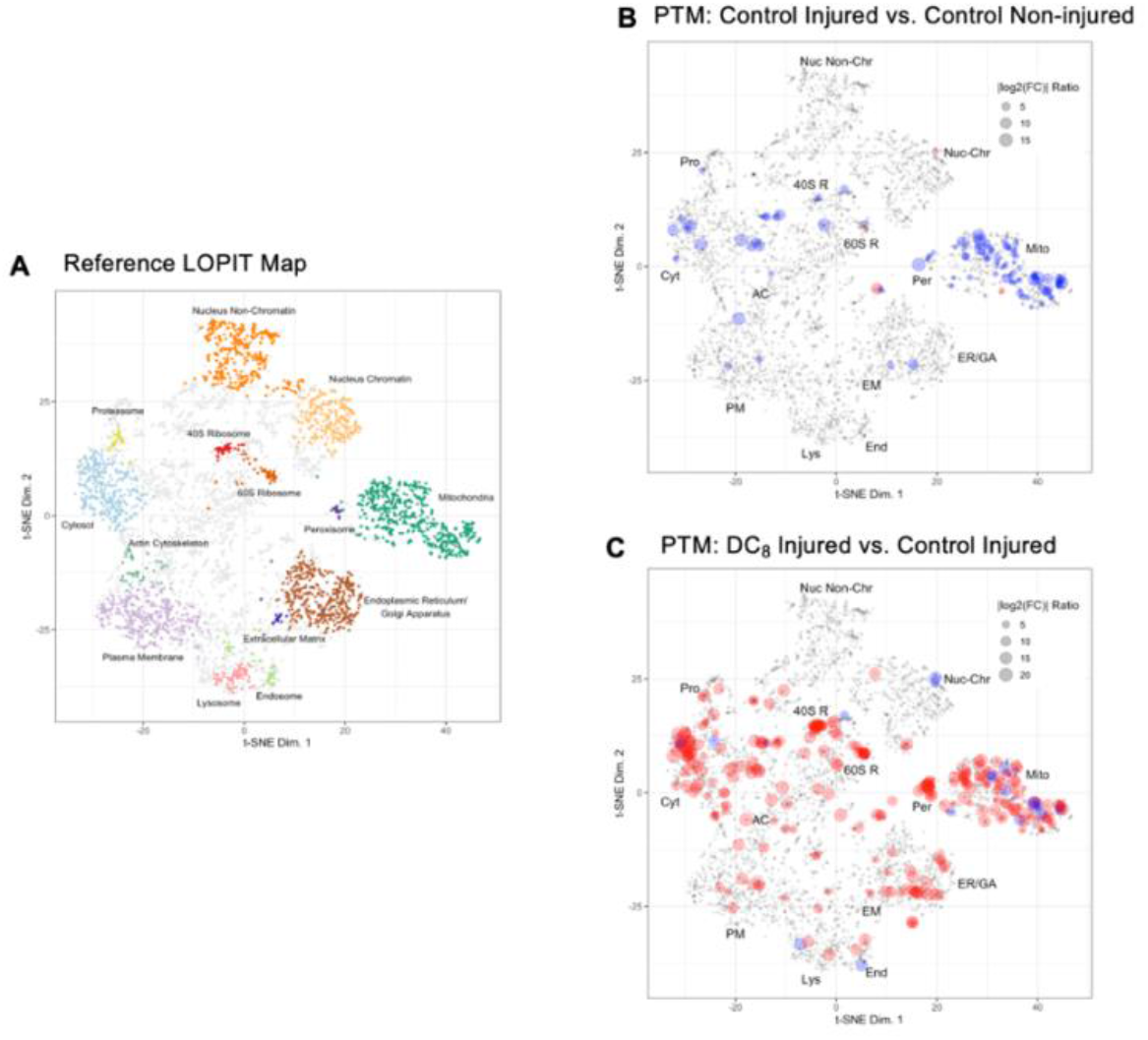
Localization of organelle proteins by isotope tagging (LOPIT) maps are a representation of protein subcellular location created by Kathryn Lilley’s lab, which are useful for analyzing proteomic data (55). They are generated by combining organelle fractionation, lysis, protein digestion, labeling with isotope tags, and quantification with mass spectrometry. Proteins are assigned to subcellular locations based on enrichment patterns, similarity to organellar markers, and co-localization with known organellar proteins (66). **A)** Distribution of LOPIT localized proteins for mouse pluripotent stem cells from (67). Assigned coordinates for 5,032 proteins in the 40S Ribosome (40S R, red), 60S Ribosome (60S R, dark orange), Actin Cytoskeleton (AC, green), Cytosol (Cyt, light blue), Endoplasmic Reticulum/Golgi Apparatus (ER/GA, brown), Endosome (End, light green), Extracellular Matrix (EM, blue), Lysosome (Lys, pink), Mitochondria (Mito, dark green), Nucleus – Chromatin (Nuc-Chr, orange), Nucleus – Non-chromatin (Nuc Non-Chr, light orange), Peroxisome (Per, purple), Plasma Membrane (PM, lavendar), Proteasome (Pro, yellow), and Undefined (grey) subcellular locations are shown. The maximum post-translational modification log_2_ Fold Change data for each protein are mapped to the LOPIT coordinates where positive fold changes are shown in red, negative fold changes are shown in blue, and dot size is indicative of the relative fold change. LOPIT maps are displayed for the following comparisons **B)** control injured vs. control non-injured comparison and **C)** DC8 injured vs. control injured comparison.

**Supplementary Figure 6:**
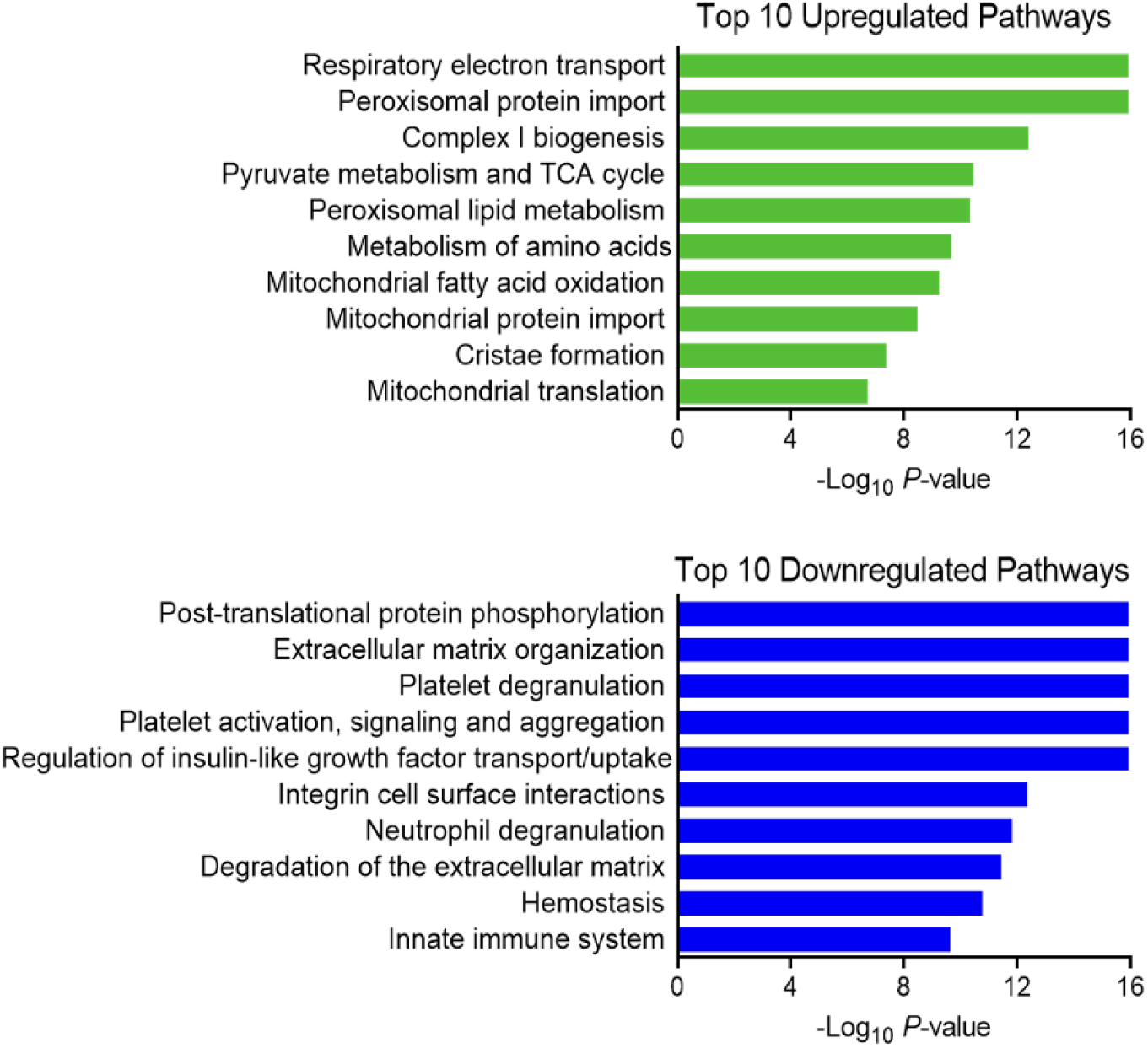
Reactome pathway analysis performed on the whole protein lysate dataset evidences the pathways that are up- and downregulated in the contralateral kidneys of DC_8_-fed animals in comparison to control-fed.

**Supplementary Figure 7:**
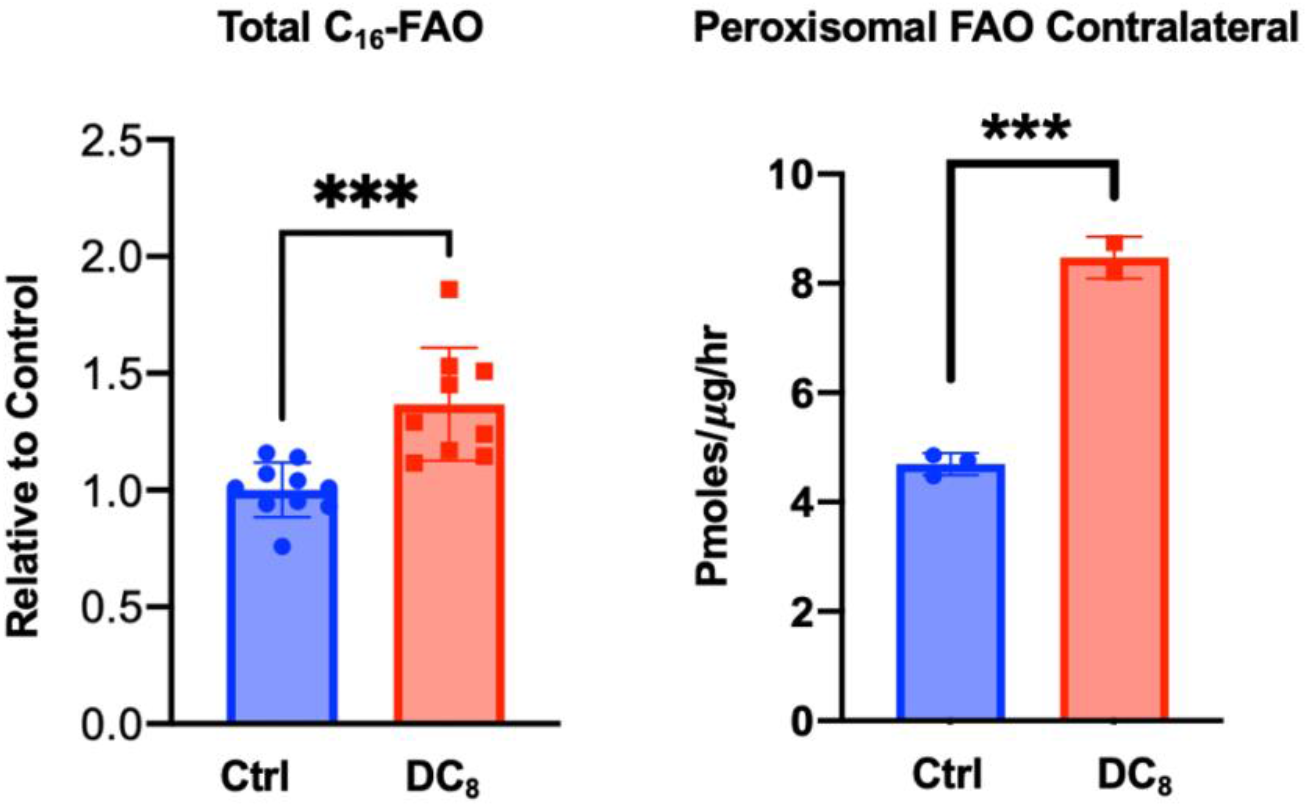
Peroxisomal oxidation of C_16_-palmitate and overall peroxisomal FAO were shown to be increased in contralateral healthy kidney homogenates from DC_8_-fed animals. N=2-9. Results are expressed as mean +- S.D. Prism 9.0.0 software (GraphPad) was used for statistical analysis. Analysis was performed using Student’s t stest. Significance was given by a p value < 0.05. *p<0.05.

## References

1. Khwaja, A., KDIGO clinical practice guidelines for acute kidney injury. Nephron Clinical Practice, 2012. 120(4): p. c179–c184.

2. Basile, D.P., M.D. Anderson, and T.A. Sutton, Pathophysiology of acute kidney injury. Compr Physiol, 2012. 2(2): p. 1303–53.

3. Kang, H.M., et al., Defective fatty acid oxidation in renal tubular epithelial cells has a key role in kidney fibrosis development. Nature medicine, 2015. 21(1): p. 37–46.

4. Menezes, L.F., et al., Fatty acid oxidation is impaired in an orthologous mouse model of autosomal dominant polycystic kidney disease. EBioMedicine, 2016. 5: p. 183–192.

5. Feingold, K.R., et al., LPS decreases fatty acid oxidation and nuclear hormone receptors in the kidney. Journal of lipid research, 2008. 49(10): p. 2179–2187.

6. Ruidera, E., et al., Fatty acid metabolism in renal ischemia. Lipids, 1988. 23(9): p. 882–4.

7. Chiba, T., et al., Sirtuin 5 Regulates Proximal Tubule Fatty Acid Oxidation to Protect against AKI. J Am Soc Nephrol, 2019. 30(12): p. 2384–2398.

8. Zhang, X., et al., Fasting induces hepatic lipid accumulation by stimulating peroxisomal dicarboxylic acid oxidation. J Biol Chem, 2021: p. 100622.

9. Westin, M.A., M.C. Hunt, and S.E. Alexson, The identification of a succinyl-CoA thioesterase suggests a novel pathway for succinate production in peroxisomes. J Biol Chem, 2005. 280(46): p. 38125–32.

10. Jin, Z., et al., Compartmentation of Metabolism of the C12-, C9-, and C5-n-dicarboxylates in Rat Liver, Investigated by Mass Isotopomer Analysis: ANAPLEROSIS FROM DODECANEDIOATE. J Biol Chem, 2015. 290(30): p. 18671–7.

11. Tserng, K.Y. and S.J. Jin, Metabolic conversion of dicarboxylic acids to succinate in rat liver homogenates. A stable isotope tracer study. J Biol Chem, 1991. 266(5): p. 2924–9.

12. Jakobs, B.S. and R.J. Wanders, Fatty acid β-oxidation in peroxisomes and mitochondria: the first, unequivocal evidence for the involvement of carnitine in shuttling propionyl-CoA from peroxisomes to mitochondria. Biochemical and biophysical research communications, 1995. 213(3): p. 1035–1041.

13. Verhoeven, N.M., et al., Phytanic acid and pristanic acid are oxidized by sequential peroxisomal and mitochondrial reactions in cultured fibroblasts. Journal of lipid research, 1998. 39(1): p. 66–74.

14. Wagner, G.R., et al., A class of reactive acyl-CoA species reveals the non-enzymatic origins of protein acylation. Cell metabolism, 2017. 25(4): p. 823–837. e8.

15. Rardin, M.J., et al., SIRT5 regulates the mitochondrial lysine succinylome and metabolic networks. Cell metabolism, 2013. 18(6): p. 920–933.

16. Park, J., et al., SIRT5-mediated lysine desuccinylation impacts diverse metabolic pathways. Molecular cell, 2013. 50(6): p. 919–930.

17. Houten, S.M., R.J. Wanders, and P. Ranea-Robles, Metabolic interactions between peroxisomes and mitochondria with a special focus on acylcarnitine metabolism. Biochimica et Biophysica Acta (BBA)-Molecular Basis of Disease, 2020. 1866(5): p. 165720.

18. Portilla, D., et al., Etomoxir-induced PPARalpha-modulated enzymes protect during acute renal failure. Am J Physiol Renal Physiol, 2000. 278(4): p. F667–75.

19. Holubarsch, C.J., et al., A double-blind randomized multicentre clinical trial to evaluate the efficacy and safety of two doses of etomoxir in comparison with placebo in patients with moderate congestive heart failure: the ERGO (etomoxir for the recovery of glucose oxidation) study. Clinical science, 2007. 113(4): p. 205–212.

20. Kovacs, W.J., et al., Localization of the pre-squalene segment of the isoprenoid biosynthetic pathway in mammalian peroxisomes. Histochemistry and cell biology, 2007. 127: p. 273–290.

21. Reszko, A.E., et al., Peroxisomal fatty acid oxidation is a substantial source of the acetyl moiety of malonyl-CoA in rat heart. Journal of Biological Chemistry, 2004. 279(19): p. 19574–19579.

22. Chiba, T., et al., Endothelial-Derived miR-17∼ 92 Promotes Angiogenesis to Protect against Renal Ischemia-Reperfusion Injury. Journal of the American Society of Nephrology, 2021. 32(3): p. 553–562.

23. Skrypnyk, N.I., R.C. Harris, and M.P. de Caestecker, Ischemia-reperfusion model of acute kidney injury and post injury fibrosis in mice. J Vis Exp, 2013(78): p. e50495–e50495.

24. Loo, D.T., TUNEL assay. In Situ Detection of DNA Damage, 2002: p. 21–30.

25. Gillet, L.C., et al., Targeted data extraction of the MS/MS spectra generated by data independent acquisition: a new concept for consistent and accurate proteome analysis. Molecular & Cellular Proteomics, 2012. 11(6).

26. Collins, B.C., et al., Multi-laboratory assessment of reproducibility, qualitative and quantitative performance of SWATH-mass spectrometry. Nature communications, 2017. 8(1): p. 291.

27. Bruderer, R., et al., Optimization of experimental parameters in data-independent mass spectrometry significantly increases depth and reproducibility of results. Molecular & Cellular Proteomics, 2017. 16(12): p. 2296–2309.

28. Bons, J., et al., In-depth analysis of the Sirtuin 5-regulated mouse brain malonylome and succinylome using library-free data-independent acquisitions. Proteomics, 2022: p. 2100371.

29. Escher, C., et al., Using i RT, a normalized retention time for more targeted measurement of peptides. Proteomics, 2012. 12(8): p. 1111–1121.

30. Storey, J.D., A direct approach to false discovery rates. Journal of the Royal Statistical Society: Series B (Statistical Methodology), 2002. 64(3): p. 479–498.

31. Burger, T., Gentle introduction to the statistical foundations of false discovery rate in quantitative proteomics. Journal of proteome research, 2018. 17(1): p. 12–22.

32. Kamburov, A., et al., ConsensusPathDB: toward a more complete picture of cell biology. Nucleic acids research, 2011. 39(suppl_1): p. D712–D717.

33. Kamburov, A., et al., ConsensusPathDB—a database for integrating human functional interaction networks. Nucleic acids research, 2009. 37(suppl_1): p. D623–D628.

34. Wickham, H. and H. Wickham, Getting started with qplot. ggplot2: elegant graphics for data analysis, 2009: p. 9–26.

35. Shannon, P., et al., Cytoscape: a software environment for integrated models of biomolecular interaction networks. Genome research, 2003. 13(11): p. 2498–2504.

36. Szklarczyk, D., et al., STRING v10: protein–protein interaction networks, integrated over the tree of life. Nucleic acids research, 2015. 43(D1): p. D447–D452.

37. Mulvey, C.M., et al., Using hyperLOPIT to perform high-resolution mapping of the spatial proteome. Nature protocols, 2017. 12(6): p. 1110–1135.

38. Gatto, L., et al., Mass-spectrometry-based spatial proteomics data analysis using pRoloc and pRolocdata. Bioinformatics, 2014. 30(9): p. 1322–1324.

39. Dunkley, T.P., et al., Localization of organelle proteins by isotope tagging (LOPIT). Molecular & Cellular Proteomics, 2004. 3(11): p. 1128–1134.

40. Christoforou, A., et al., A draft map of the mouse pluripotent stem cell spatial proteome. Nature communications, 2016. 7(1): p. 9992.

41. Van der Maaten, L. and G. Hinton, Visualizing data using t-SNE. Journal of machine learning research, 2008. 9(11).

42. Burton, J.B., et al., Pattern analysis of organellar maps for interpretation of proteomic data. Proteomes, 2022. 10(2): p. 18.

43. Zhang, Y., et al., Lysine desuccinylase SIRT5 binds to cardiolipin and regulates the electron transport chain. Journal of Biological Chemistry, 2017. 292(24): p. 10239–10249.

44. Bharathi, S.S., et al., Role of mitochondrial acyl-CoA dehydrogenases in the metabolism of dicarboxylic fatty acids. Biochemical and biophysical research communications, 2020. 527(1): p. 162–166.

45. Grego, A. and G. Mingrone, Dicarboxylic acids, an alternate fuel substrate in parenteral nutrition: an update. Clinical Nutrition, 1995. 14(3): p. 143–148.

46. Membrez, M., et al., Six weeks’ sebacic acid supplementation improves fasting plasma glucose, HbA1c and glucose tolerance in db/db mice. Diabetes, Obesity and Metabolism, 2010. 12(12): p. 1120–1126.

47. Goeckermann, J.A. and E.L. Vigil, Peroxisome development in the metanephric kidney of mouse. J Histochem Cytochem, 1975. 23(12): p. 957–73.

48. Litwin, J.A., et al., Immunocytochemical demonstration of peroxisomal enzymes in human kidney biopsies. Virchows Arch B Cell Pathol Incl Mol Pathol, 1988. 54(4): p. 207–13.

49. Litwin, J.A., et al., Detection of peroxisomes in human liver and kidney fixed with formalin and embedded in paraffin: the use of catalase and lipid beta-oxidation enzymes as immunocytochemical markers. Histochem J, 1988. 20(3): p. 165–73.

50. Vasko, R., Peroxisomes and Kidney Injury. Antioxid Redox Signal, 2016. 25(4): p. 217–31.

51. Gulati, S., et al., Alterations of peroxisomal function in ischemia-reperfusion injury of rat kidney. Biochim Biophys Acta, 1993. 1182(3): p. 291–8.

52. Singh, A.K. and S. Gulati, Morphology of peroxisomes after ischemia-reperfusion injury. Ann N Y Acad Sci, 1994. 723: p. 403–5.

53. Hasegawa, K., et al., Kidney-specific overexpression of Sirt1 protects against acute kidney injury by retaining peroxisome function. J Biol Chem, 2010. 285(17): p. 13045–56.

54. Rodriguez-Calvo, R., et al., AICAR Protects against High Palmitate/High Insulin-Induced Intramyocellular Lipid Accumulation and Insulin Resistance in HL-1 Cardiac Cells by Inducing PPAR-Target Gene Expression. PPAR Res, 2015. 2015: p. 785783.

55. Huang, H., et al., Inhibitors of Fatty Acid Synthesis Induce PPAR alpha -Regulated Fatty Acid beta -Oxidative Genes: Synergistic Roles of L-FABP and Glucose. PPAR Res, 2013. 2013: p. 865604.

56. Tsukamoto, T., et al., Vaticanol C, a resveratrol tetramer, activates PPARalpha and PPARbeta/delta in vitro and in vivo. Nutr Metab (Lond), 2010. 7: p. 46.

57. Tan, L., J.T. Yu, and H.S. Guan, Resveratrol exerts pharmacological preconditioning by activating PGC-1alpha. Med Hypotheses, 2008. 71(5): p. 664–7.

58. Planavila, A., et al., Sirt1 acts in association with PPARalpha to protect the heart from hypertrophy, metabolic dysregulation, and inflammation. Cardiovasc Res, 2011. 90(2): p. 276–84.

59. Abdel-Razek, E.A.-N., A.M. Abo-Youssef, and A.A. Azouz, Benzbromarone mitigates cisplatin nephrotoxicity involving enhanced peroxisome proliferator-activated receptor alpha (PPAR-α) expression. Life sciences, 2020. 243: p. 117272.

60. Kim, S.-J., et al., Protective roles of fenofibrate against cisplatin-induced ototoxicity by the rescue of peroxisomal and mitochondrial dysfunction. Toxicology and Applied Pharmacology, 2018. 353: p. 43–54.

61. Litwin, J., et al., Immunocytochemical demonstration of peroxisomal enzymes in human kidney biopsies. Virchows Archiv B, 1987. 54(1): p. 207–213.

62. Litwin, J., et al., Detection of peroxisomes in human liver and kidney fixed with formalin and embedded in paraffin: the use of catalase and lipid β-oxidation enzymes as immunocytochemical markers. The Histochemical Journal, 1988. 20(3): p. 165–173.

63. dos Santos, N.A.G., et al., Cisplatin-induced nephrotoxicity and targets of nephroprotection: an update. Archives of toxicology, 2012. 86(8): p. 1233–1250.

64. Pabla, N. and Z. Dong, Cisplatin nephrotoxicity: mechanisms and renoprotective strategies. Kidney international, 2008. 73(9): p. 994–1007.

65. Peres, L.A.B. and A.D.d. Cunha Júnior, Acute nephrotoxicity of cisplatin: molecular mechanisms. Brazilian Journal of Nephrology, 2013. 35: p. 332–340.

66. Vasko, R., et al., Endothelial peroxisomal dysfunction and impaired pexophagy promotes oxidative damage in lipopolysaccharide-induced acute kidney injury. Antioxidants & redox signaling, 2013. 19(3): p. 211–230.

67. Trub, A.G. and M.D. Hirschey, Reactive Acyl-CoA Species Modify Proteins and Induce Carbon Stress. Trends Biochem Sci, 2018. 43(5): p. 369–379.

68. Wagner, G.R., et al., A Class of Reactive Acyl-CoA Species Reveals the Non-enzymatic Origins of Protein Acylation. Cell Metab, 2017. 25(4): p. 823–837 e8.

69. Tsuchiya, Y., U. Pham, and I. Gout, Methods for measuring CoA and CoA derivatives in biological samples. Biochem Soc Trans, 2014. 42(4): p. 1107–11.

