## Supplemental material for "Dicarboxylic acid supplementation protects from acute kidney injury via stimulation of renal peroxisomal activity"

### SUPPLEMENTAL APPENDIX

### Table of Contents

**Supplemental Methods** ……………………………………………………………………………..2

Proteomic Analysis …………………………………………………………………………..2

Protein digestion and desalting: Sample cohort……………………………………………..2

Protein digestion and desalting: Spectral library of the mouse kidney whole lysate …………..3

Mass Spectrometry Analysis: Sample cohort………………………………………………..4

Mass Spectrometry Analysis: Spectral Library…………………………………………...5

**Data Analysis** ………………..………………………………………………………………….5

Spectral library generation ………………………………………………………………….5

DIA Data Processing and Statistical Analysis………………………………………..……..……..6

Spectral library generation ……………………………………………………………………..5

Enrichment analysis …………………………………………………………………………..7

Subcellular localization……………………………………..……………………………...………..7

LOPIT Organellar Protein Distribution……………………………………………………….…..7

**Supplementary figures** ……………………………………………………………………….8

Supplementary figure 1…………………………………………………………………………..8

Supplementary figure 2…………………………………………………………………………..9

Supplementary figure 3…………………………………………………………………...…..10

Supplementary figure 4……………………………………………………………………….11

**Supplementary Figures**

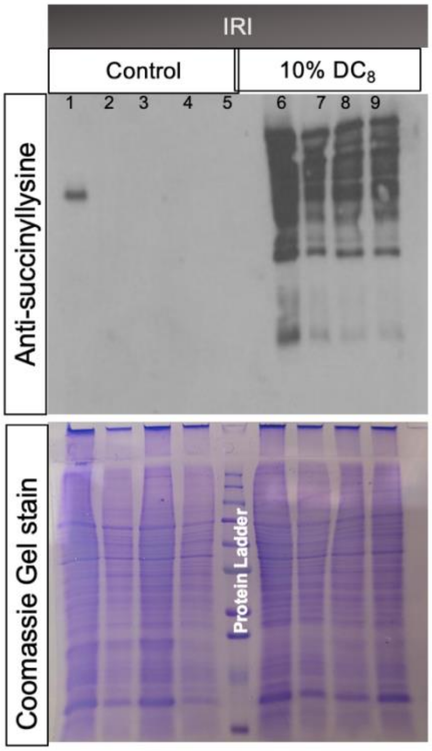


**Supplementary Figure 1:** Kidney succinylation is increased in $DC_8$-fed animals, both in contralateral healthy and in kidneys that were subjected to IRI. N=4.

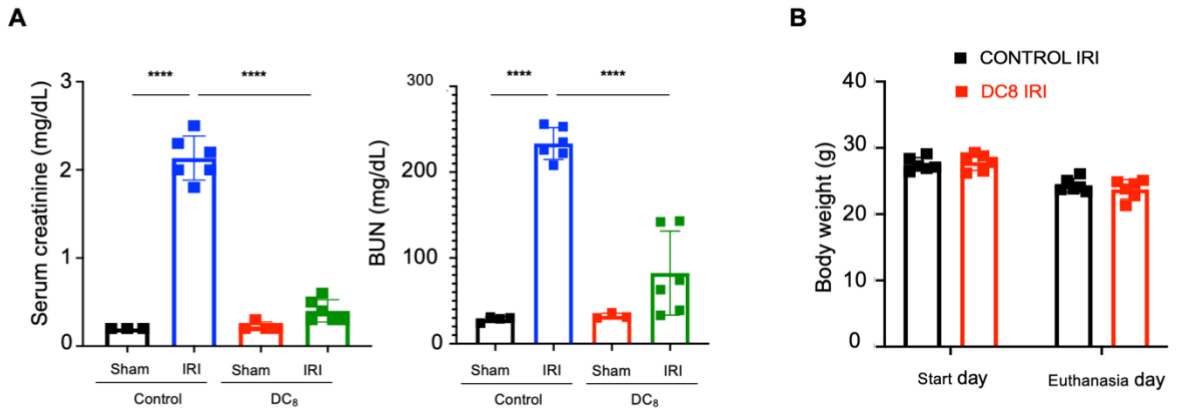


**Supplementary Figure 2:** Comparison between Sham and IRI animals fed with either a control- or $DC_8$-diet. Creatinine and BUN show no significant differences between control- and $DC_8$-fed animals that were solely subjected to a unilateral nephrectomy 24 hours prior to sacrifice, whereas when mice undergo IRI, $DC_8$-fed are significantly protected from the injury. N= 6 (IRI) and 3 (sham). Results are expressed as mean +- S.D. Prism 9.0.0 software (GraphPad) was used for statistical analysis. Analysis was performed using Student's t test. Significance was given by a p value < 0.05. *p<0.05 ****p< 0.0001.

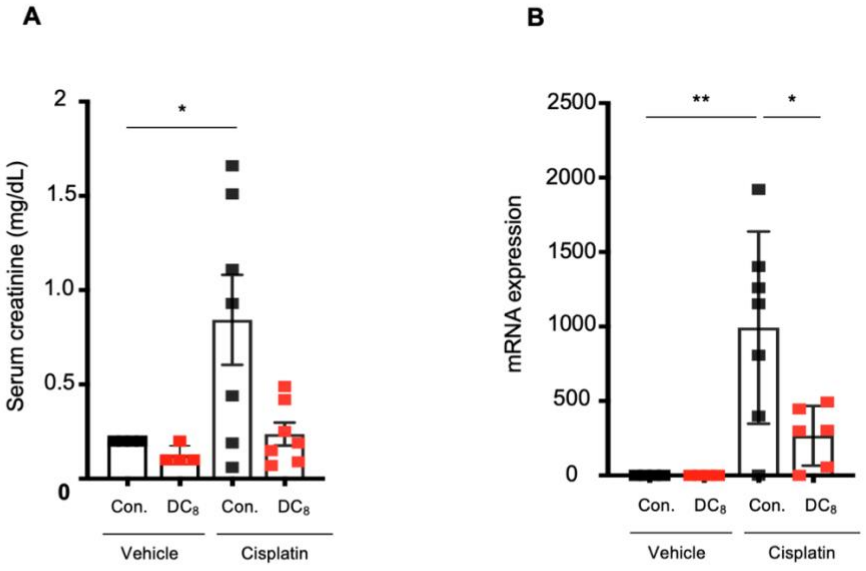


**Supplementary Figure 3:** $DC_8$-fed animals are protected from Cisplatin-induced AKI. A) Serum creatinine and renal NGAL expression; B) H&E staining. N=7 (cisplatin groups) and 4 (vehicle groups). Results are expressed as mean +- S.D. Prism 9.0.0 software (GraphPad) was used for statistical analysis. Analysis was performed using Student's t test. Significance was given by a p value < 0.05. *p<0.05.

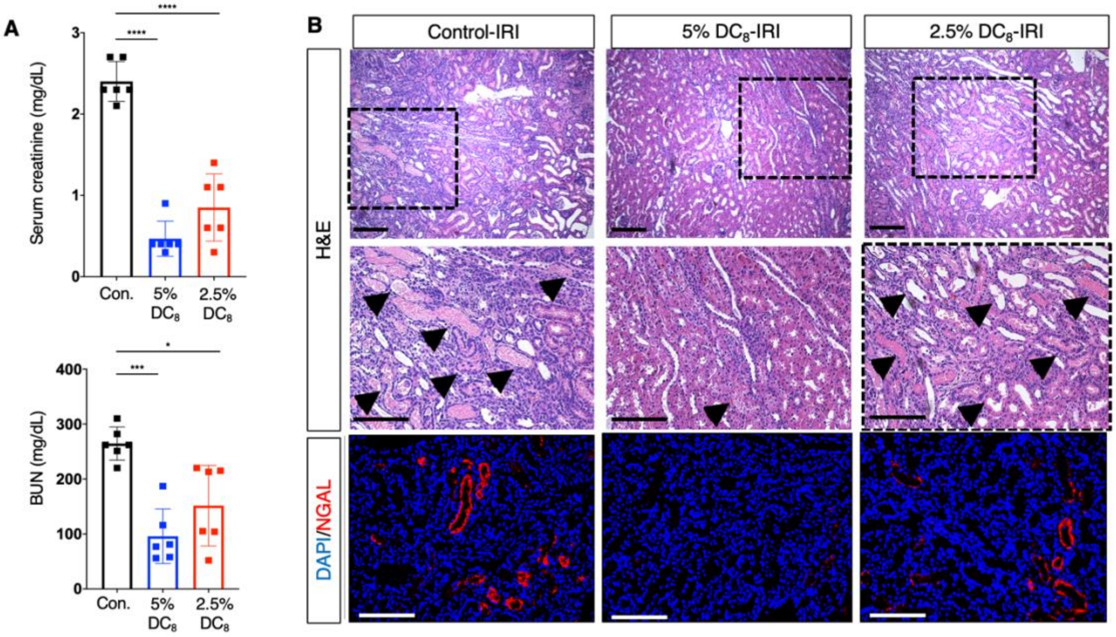


**Supplementary Figure 4:** $DC_8$ protects animals from AKI at a concentration as low as 5% w/w. A) Creatinine and BUN; B) H&E staining and immunofluorescence for NGAL and DAPI. N=6. Scale bars represent a 10x and 20x magnification, respectively. Results are expressed as mean +- S.D. Prism 9.0.0 software (GraphPad) was used for statistical analysis. Analysis was performed using Student's t test. Significance was given by a p value < 0.05. *p<0.05 **p<0.01 ***p<0.001 ****p<0.0001.

**A** Mitochondrial and peroxisomal succinylated protein groups

| | All quantified proteins | Down-reg. in $DC_8$ vs Control, injured | Up-reg. in $DC_8$ vs Control, injured | Down-reg. in $DC_8$ vs Control, non-injured | Up-reg. in $DC_8$ vs Control, non-injured |
|---|---|---|---|---|---|
| **Total** | 1,160 | 95 | 657 | 262 | 461 |
| **Mitochondrion** | 348 | 58 | 195 | 179 | 103 |
| **Peroxisome** | 46 | 2 | 44 | 4 | 38 |

**B** All quantifiable succinylated protein groups

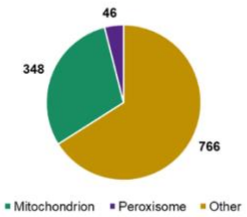


**Supplementary Figure 5:** Lysine succinylome-based mass spectrometry evidences that $DC_8$ induces a drastic remodeling of mitochondrial and peroxisomal succinylomes in both injured and non-injured kidneys. Protein subcellular localization information was retrieved using stringAPP (Szklarczyk *et al.*, 2015, *Nucleic Acids Res*, 43, D447-52), and the compartment score threshold was set at 4.5. 'Total' in **A** and 'Other' in **B** include proteins not assigned to mitochondrial nor peroxisomal compartments as well as proteins for which no subcellular localization information was retrieved. **A)** Summary of all succinylated proteins as well as mitochondrial and peroxisomal succinylated proteins that were quantified and significantly down- and up-regulated for $DC_8$ vs Control comparisons in both injured and non-injured kidneys. **B)** Proportion of proteins assigned to mitochondrion (green), peroxisome (purple) and other organelles (beige) among all quantified succinylated protein groups.

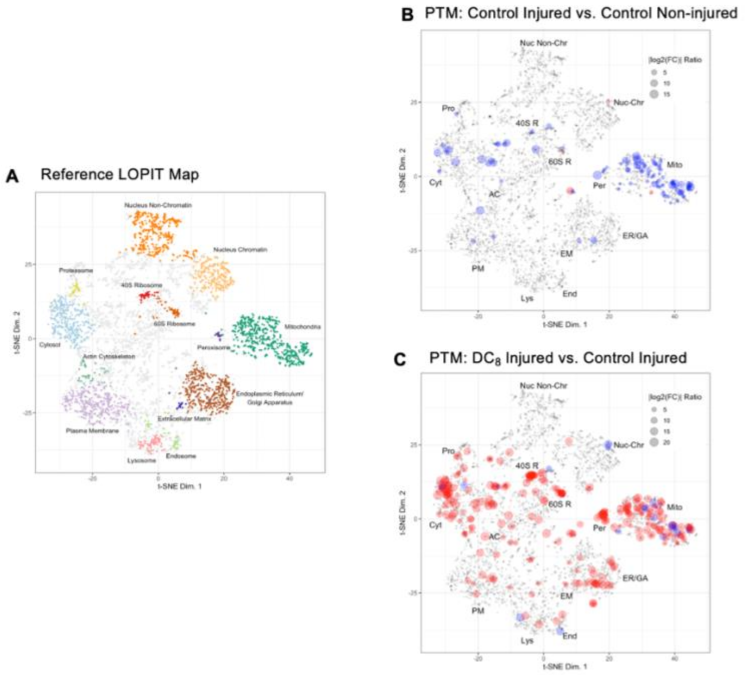


**Supplementary Figure 6:** Localization of organelle proteins by isotope tagging (LOPIT) maps are a representation of protein subcellular location created by Kathryn Lilley's lab, which are useful for analyzing proteomic data (55). They are generated by combining organelle fractionation, lysis, protein digestion, labeling with isotope tags, and quantification with mass spectrometry. Proteins are assigned to subcellular locations based on enrichment patterns, similarity to organellar markers, and co-localization with known organellar proteins (66). **A)** Distribution of LOPIT localized proteins for mouse pluripotent stem cells from (67). Assigned coordinates for 5,032 proteins in the 40S Ribosome (40S R, red), 60S Ribosome (60S R, dark orange), Actin Cytoskeleton (AC, green), Cytosol (Cyt, light blue), Endoplasmic Reticulum/Golgi Apparatus (ER/GA, brown), Endosome (End, light green), Extracellular Matrix (EM, blue), Lysosome (Lys, pink), Mitochondria (Mito, dark green), Nucleus – Chromatin (Nuc-Chr, orange), Nucleus – Non-chromatin (Nuc Non-Chr, light orange), Peroxisome (Per, purple), Plasma Membrane (PM, lavendar), Proteasome (Pro, yellow), and Undefined (grey) subcellular locations are shown. The maximum post-translational modification $\log_2$ Fold Change data for each protein are mapped to the LOPIT coordinates where positive fold changes are shown in red, negative fold changes are shown in blue, and dot size is indicative of the relative fold change. LOPIT maps are displayed for the following comparisons **B)** control injured vs. control non-injured comparison and **C)** DC8 injured vs. control injured comparison.

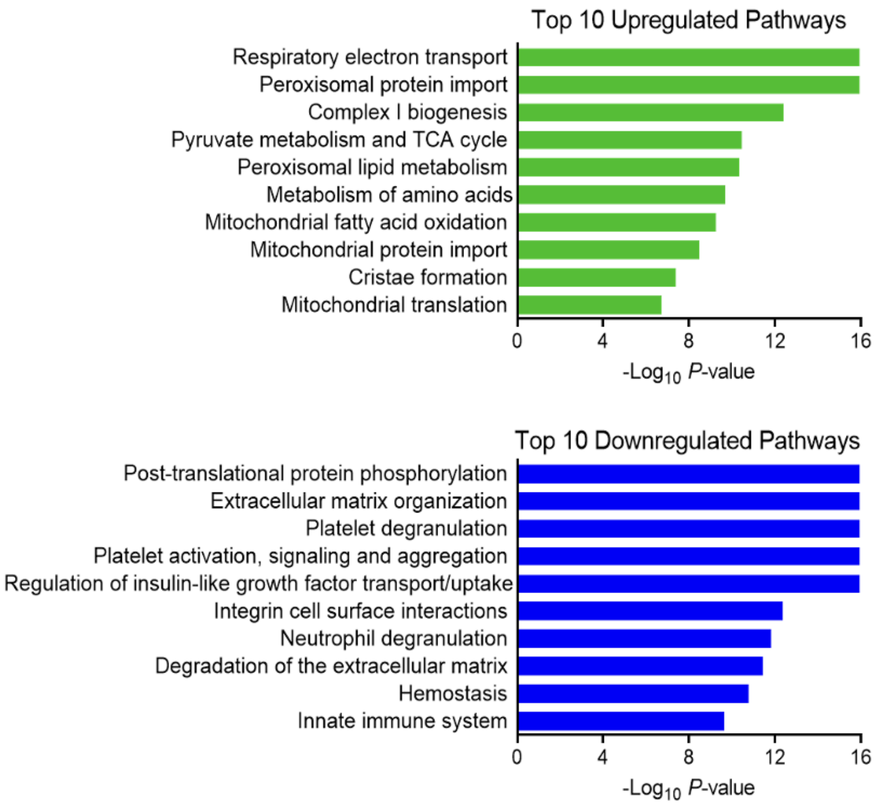


**Supplementary Figure 7:** Reactome pathway analysis performed on the whole protein lysate dataset evidences the pathways that are up- and downregulated in the contralateral kidneys of $DC_8$-fed animals in comparison to control-fed.

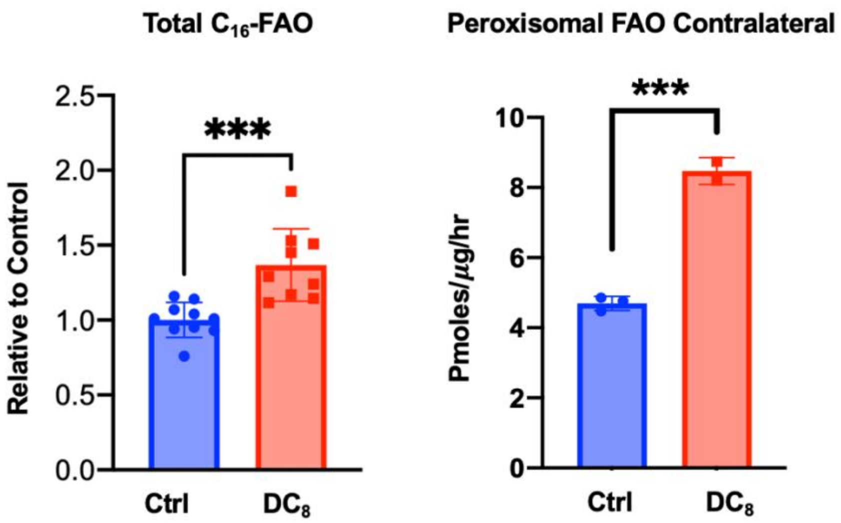


**Supplementary Figure 8:** Peroxisomal oxidation of $C_{16}$-palmitate and overall peroxisomal FAO were shown to be increased in contralateral healthy kidney homogenates from $DC_8$-fed animals. N=2-9. Results are expressed as mean +- S.D. Prism 9.0.0 software (GraphPad) was used for statistical analysis. Analysis was performed using Student's t test. Significance was given by a p value < 0.05. *p<0.05.
